# Protein prenylation and Hsp40 in thermotolerance of *Plasmodium falciparum* malaria parasites

**DOI:** 10.1101/842468

**Authors:** Emily S. Mathews, Andrew J. Jezewski, Audrey R. Odom John

## Abstract

During its complex life cycle, the malaria parasite survives dramatic changes in environmental temperature. Protein prenylation is required during asexual replication of *Plasmodium falciparum*, and heat shock protein 40 (HSP40; PF3D7_1437900) is post-translationally modified with a 15-carbon farnesyl isoprenyl group. In other organisms, farnesylation of Hsp40 orthologs controls its localization and function, including temperature stress survival. In this work, we find that plastidial isopentenyl pyrophosphate (IPP) synthesis and protein farnesylation are required for malaria parasite survival after cold and heat shock. Furthermore, loss of HSP40 farnesylation alters its membrane attachment and interaction with proteins involved in crucial biological processes, such as glycolysis and cytoskeletal organization. Together, this work reveals that farnesylation of HSP40 in *P. falciparum* is a novel essential function of plastidial isoprenoid biosynthesis. We propose a model by which farnesyl-HSP40 promotes parasite thermotolerance and facilitates vesicular trafficking through its interaction with client proteins.

## Introduction

Infection with the protozoan parasite *Plasmodium falciparum* causes the majority of cases of severe and fatal malaria. *P. falciparum* must recognize and adapt to dramatic temperature swings, as its complex life cycle requires development in both an invertebrate mosquito vector and the warm-blooded vertebrate human host. Temperature is critical at every stage of the parasite life cycle. In the mosquito vector, many temperature-sensitive factors contribute to human transmission, such as biting rate, vector longevity, parasite development, and vector competence^1^. Human infection begins upon the bloodmeal of a female *Anopheles* mosquito. Entering the human host, where the normal physiological temperature is 37°C, sporozoite-stage parasites experience heat shock. However, this temperature stress is necessary for efficient hepatocyte infection and the resulting amplification of infection^2, 3^. The parasite emerges from the liver to initiate asexual replication within erythrocytes, the clinically symptomatic stage of *Plasmodium* infection. A pathognomonic feature of falciparum malaria is periodic episodes of fever (to 41°C or more) recurring every 48 hours, corresponding to the synchronous rupture of infected erythrocytes and daughter merozoite release^4^. In contrast, the sexual-stage parasites that return to the mosquito vector are again exposed to cold temperature shock, as the parasite must now re-adjust to approximately 25°C. While temperature fluctuations are an inherent part of the malaria life cycle, how the parasite copes with thermal stress is not well understood.

Temperature also regulates malaria parasite virulence and antimalarial sensitivity. Controlled hypothermia (32°C) has been clinically used to improve outcomes of severe cerebral malaria^5^. *In vitro*, hypothermia (32°C) inhibits *P. falciparum* growth,^6^ and a similar effect occurs at lower temperatures (28°C)^7^. While the potency of many antimalarials (e.g., chloroquine, mefloquine, and pyronaridine) are unaffected by lower temperatures^6, 8^, susceptibility to artemisinin—the backbone of front-line artemisinin-based combination therapies—is modulated by both cold and heat stresses^6, 8^. As temperature fluctuations are an inherent part of the *P. falciparum* life cycle, this common environmental stress may impact the ability of antimalarials to influence essential parasite targets. Rising rates of delayed clearance to artemisinin-based combination therapies, including resistance to artemisinin partner drugs, has raised concerns about emerging multi-drug resistance^9–17^. Thus, there is a pressing need to identify essential survival pathways in *P. falciparum*, such as thermotolerance, in order to support ongoing development of new antimalarial agents.

During intraerythrocytic development, *P. falciparum* assembles isoprenoids *de novo* through the methylerythritol 4-phosphate (MEP) pathway^18–20^, localized to the unusual plastidial organelle of the parasite, the apicoplast. Chemical inhibition of this pathway by the small molecule fosmidomycin (FSM) is lethal to malaria parasites^18^. FSM-mediated growth inhibition can be rescued by supplementation with isoprenoids such as isopentenyl pyrophosphate (IPP)^21, 22^. These studies validate the essentiality of isoprenoid synthesis in asexual *P. falciparum*, but there have been long-standing questions about which biological processes in the parasite require apicoplast isoprenoid biosynthesis. Protein prenylation appears to be one core essential function of isoprenoid biosynthesis in malaria parasites^23–27^. During protein prenylation, either a farnesyl (FPP; 15-carbon) or geranylgeranyl (GGPP; 20-carbon) isoprenyl group is post-translationally attached to C-terminal cysteines by one of three well-characterized prenyltransferases, farnesyltransferase (FTase) and geranylgeranyltransferases type I and type II (GGTase I, GGTase II). Chemical inhibition of prenyltransferases with small molecules (e.g., FTase inhibitor FTI-277) inhibits parasite growth^24–29^, providing compelling evidence that prenylated malarial proteins and their unidentified downstream biological processes include potential antimalarial targets. We and others have used chemical labeling to characterize the complete prenylated proteome of intraerythrocytic *P. falciparum*^30, 31^. These studies identify a single heat shock protein 40 (HSP40; PF3D7_1437900) as robustly farnesylated during intraerythrocytic replication.

Heat shock proteins are necessary for protein folding and stabilization. Importantly, heat shock proteins play a vital role in surviving cellular insults that might otherwise be lethal, and therefore heat shock protein expression is upregulated under diverse cellular stresses, including heat and cold shock. The main functions of Hsp40 family members are to identify and bind partially misfolded proteins, in order to initiate Hsp70-mediated refolding. As heat shock proteins have a known role in temperature-dependent survival, it is perhaps unsurprising that roughly 2% of the *P. falciparum* genome is dedicated to molecular chaperones, including a large number of heat shock proteins^32^. HSP40 is a member of an expanded Hsp40 family in *P. falciparum* comprising 49 total members^33^. The majority of Hsp40 family members in *P. falciparum* are unique and not shared with other Apicomplexa^33, 34^. HSP40 is predicted to be the only canonical Hsp40 and the main co-chaperone of Hsp70 in *P. falciparum* because of its similar heat inducibility and localization^35^. The lack of additional canonical Hsp40s in *P. falciparum* suggests that HSP40 is necessary for parasite development. Importantly, HSP40 is the only prenylated heat shock protein in *P. falciparum*^30, 31^. In yeast, prenylation of the HSP40 homolog YDJ1 is required for thermotolerance, protein localization, and interaction with client proteins^36–38^. In *P. falciparum,* the role of farnesylated HSP40 (farnesyl-HSP40) has not previously been investigated.

In this study, we investigate the role of farnesylation on the function of HSP40 during intraerythrocytic growth of *P. falciparum*. We find that farnesylation, but not geranylgeranylation, contributes to parasite survival following either heat (40°C) or cold shock (25°C). We demonstrate that HSP40 is likely essential, requires farnesylation for membrane association and for interaction with proteins in essential pathways in the parasite. We thus provide the first evidence that farnesylation of HSP40 influences its function in *P. falciparum*. In addition, our work suggests that loss of HSP40 prenylation contributes to parasite death that results from reduced apicoplast function or inhibition of protein prenylation, and also confers a hypersensitivity to temperature stress.

## Results

### Thermotolerance in malaria parasites requires IPP synthesis and protein farnesylation

The malaria parasite must adapt to diverse conditions, such as temperature stress, throughout its complex life cycle. Heat shock proteins play an important role in the ability of the parasite to survive temperature stress^39–42^. Because *P. falciparum* HSP40 is post-translationally farnesylated, we hypothesized that farnesylation of HSP40 is required for growth during temperature stress. We tested this hypothesis by inhibiting protein prenylation and applying either heat or cold stress. Chemically diverse small molecule inhibitors impact protein prenylation during asexual replication of *Plasmodium* spp. For example, treatment with FSM, which inhibits upstream isoprenoid biosynthesis and therefore synthesis of prenylphosphates, reduces downstream protein prenylation^23^. Well-validated prenyltransferase inhibitors, such as FTI-277 (FTI), BMS-388891 (BMS), and GGTI-298 (GGTI), directly reduce levels of protein prenylation in *P. falciparum*^28, 43^ (Fig. 1A). We tested whether inhibition of prenylphosphate synthesis or prenyltransferases influenced parasite growth following heat (40°C) or cold (25°C) stress. These temperatures were selected to emulate temperatures in which the parasite is exposed during febrile episodes (for heat stress) and during transmission to the mosquito vector (for cold stress). Parasites were pre-treated with inhibitor 24 hours prior to temperature shock. Parasite growth was evaluated by flow cytometry (Fig. S1) for 6 days post-shock (Fig. 1B).

While untreated parasites readily recover following brief heat or cold shock (Fig. 1C, D), we find that parasite survival under heat (Fig. 1C) or cold (Fig. 1D) stress was significantly attenuated upon non-lethal inhibition of IPP synthesis by FSM. FTI and BMS are chemically distinct, well validated protein farnesyltransferase inhibitors, while GGTI inhibits protein geranylgeranylation. Using these inhibitors, we find that chemical inhibition of protein farnesylation, but not geranylgeranylation, significantly impairs temperature stress recovery (Fig. 1E; Fig. S2).

Taking advantage of the fact that FSM inhibits production of all isoprenoid products downstream of IPP, we employed chemical supplementation in order to determine which isoprenoids are required for temperature stress survival in malaria parasites. We find that supplementation with IPP or farnesol (F-OL), a 15-carbon farnesyl alcohol, rescues FSM-treated parasites after both heat and cold stress (Fig. 2B-D, F-H). In contrast, supplementation with geranylgeraniol (GG-OL), a 20-carbon geranylgeranyl alcohol, does not rescue growth of FSM-treated parasites after temperature shock (Fig. 2E, I). Altogether, these data establish that loss of isoprenoid biosynthesis or protein farnesylation sensitize malaria parasites to changes in temperature. Protein farnesylation is thus required for parasite survival following moderate, non-lethal temperature stress, in which heat shock proteins have a canonical biological role. These data raise the possibility that HSP40—one of only 4 farnesylated proteins in *P. falciparum* and the sole farnesylated heat shock protein^30, 31^—might be vital for parasite survival.

**Figure 1.**
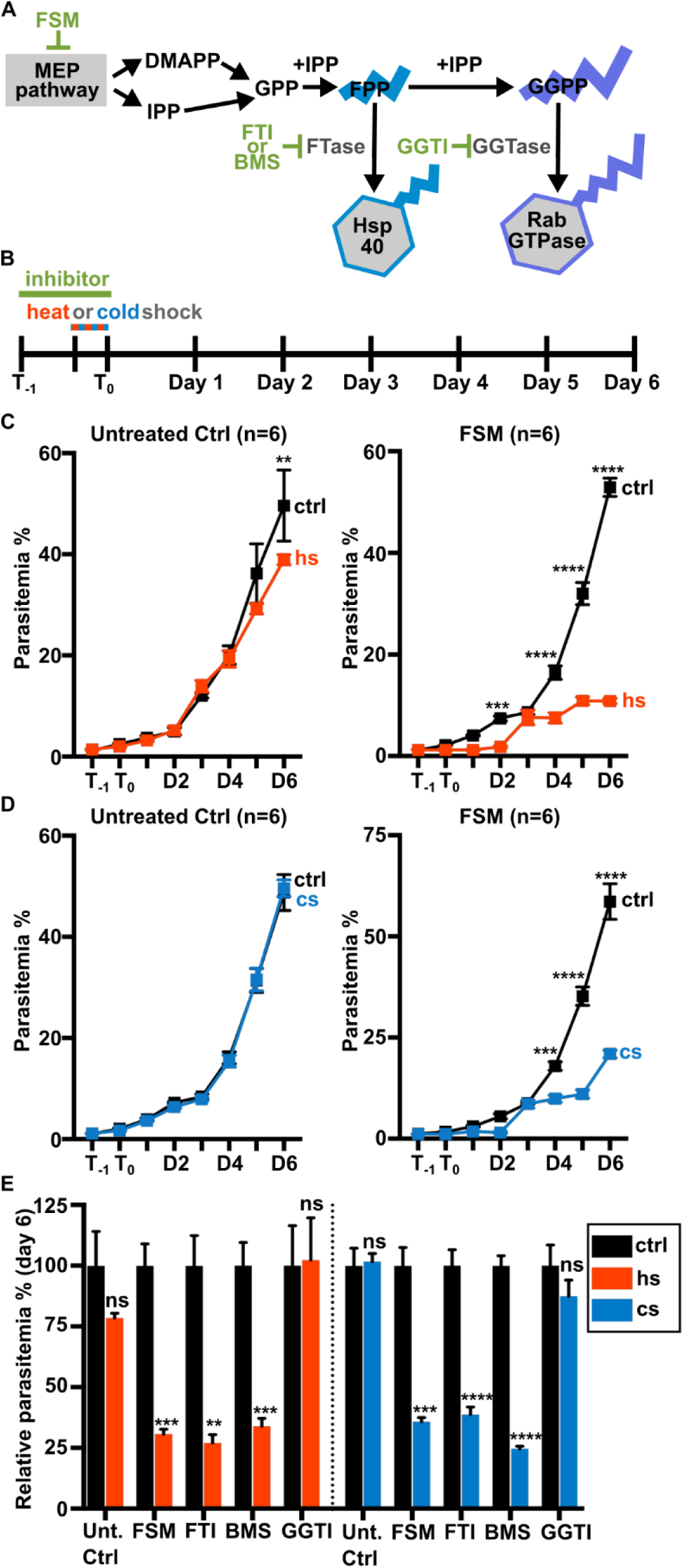
Growth under temperature stress requires IPP synthesis and protein farnesylation. (A) Prenylphosphate substrates for protein prenylation are derived from the non-mevalonate MEP pathway. MEP pathway products, IPP and dimethylallyl pyrophosphate (DMAPP) serve as precursors to FPP used by FTase and GGPP used by GGTase in protein prenylation. FSM treatment inhibits production of IPP and DMAPP. Farnesyltransferase inhibitors (FTI or BMS) inhibit protein farnesylation, while geranylgeranyltransferase inhibitors (GGTI) prevent protein geranylgeranylation. (B) Parasites were treated with FSM (5 μM), farnesyltransferase inhibitors [FTI (10μM) and BMS (200nM)] or geranylgeranyltransferase inhibitor GGTI (2μM) for 24 hours prior to a 6 hour heat (40°C) or cold (25°C) shock. (C,D) FSM-treated parasite growth is significantly reduced after heat shock (C) and cold shock (D). (E) Inhibition of farnesylation by treating parasites with FTI or BMS significantly reduced growth after temperature stress. Growth in GGTI-treated parasites is unchanged after heat or cold shock. (C-E) n = 6; ** p≤0.01; *** p≤0.001; **** p≤0.0001, (C,D) 2-way ANOVA, p-values adjusted for multiple comparisons using Sidak’s multiple comparison test, (E) within each treatment group the normalized control was compared to temperature shock sample by unpaired t-test with Welch’s correction. Abbreviations: control (ctrl), heat shock (hs), cold shock (cs).

**Supplemental Figure 1.**
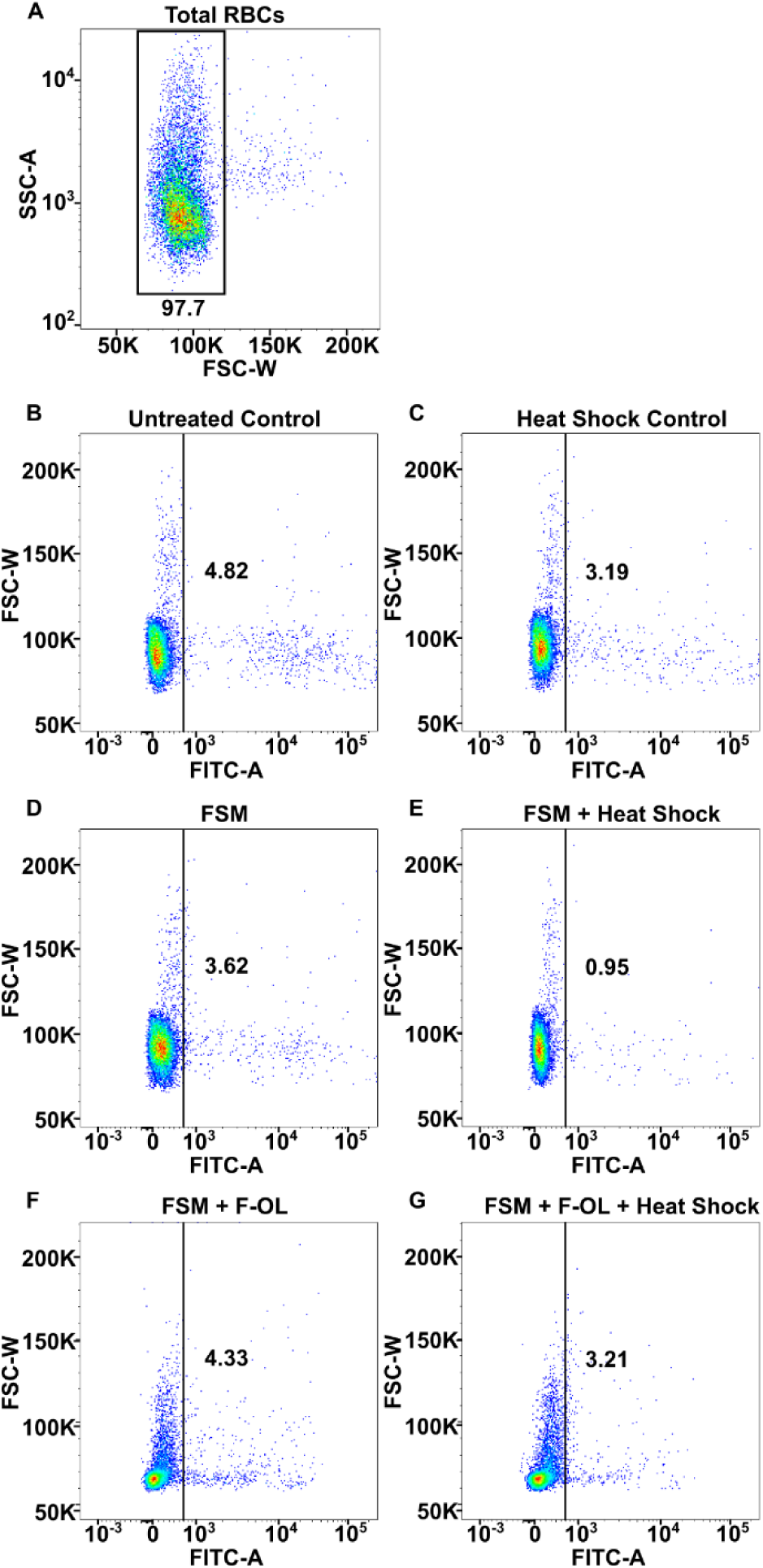
Gating strategy for infected RBCs. Uninfected erythrocytes were used as a control for gating purposes. (A) Total RBCs were gated based on the side scatter and forward scatter (SSC-A/FSC-W) profile in the dot plot. (B-G) Parasite infected RBCs were gated based on forward scatter and fluorescein isothiocyanate (FSC-W/FITC-A). Gating was kept consistent between all treatment groups. Differences in number of parasite-infected RBCs are easily visible in dot plots. For example, heat shocked FSM-treated parasites had fewer infected RBCs (E). Farnesol (F-OL) can rescue this temperature sensitive growth (G).

**Supplemental Figure 2.**
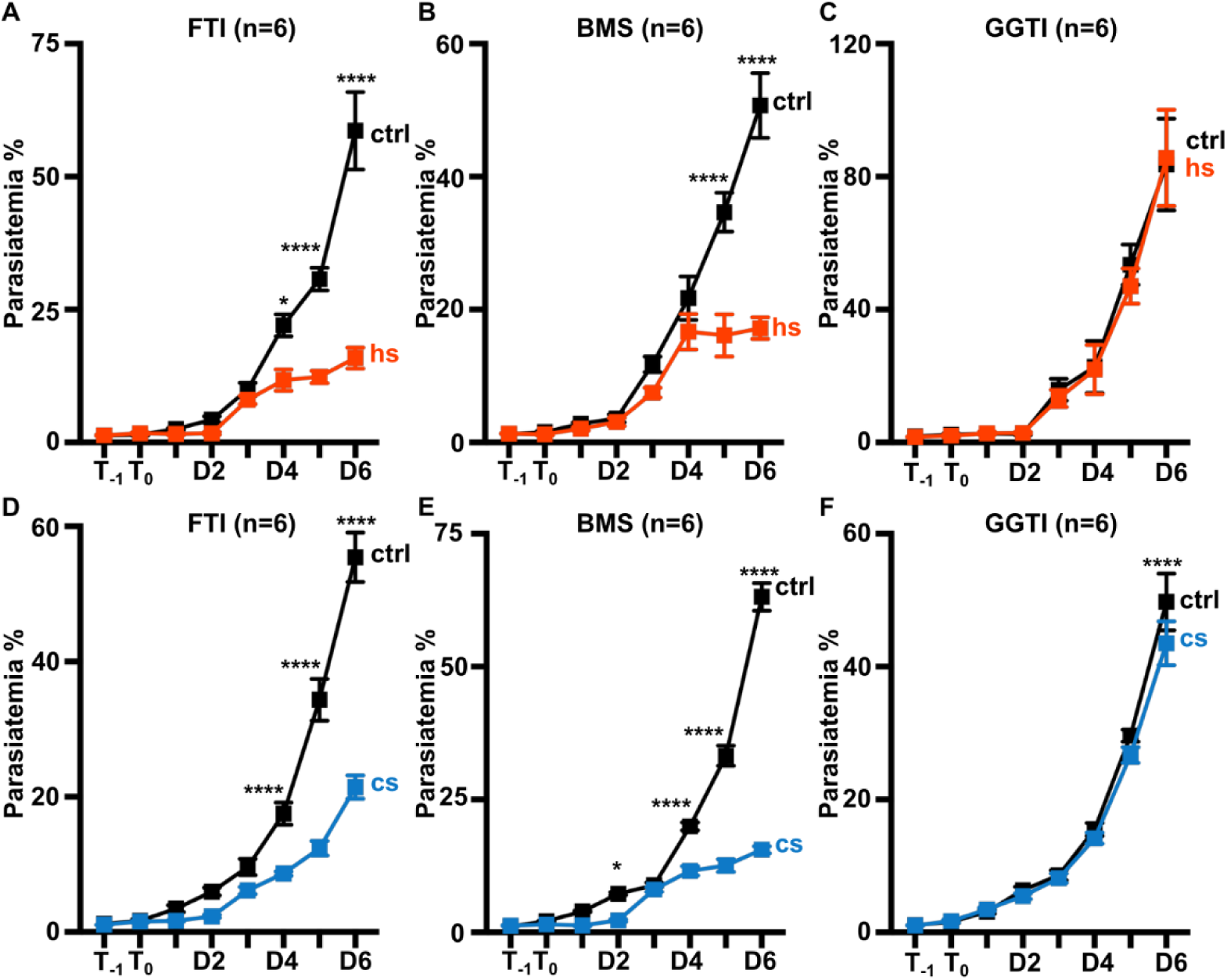
Protein farnesylation, but not geranylgeranylation, is required for thermotolerance in malaria parasites. Parasites were treated with farnesyltransferase inhibitors [FTI (10μM) and BMS (200nM)] or geranylgeranyltransferase inhibitor GGTI (2μM) for 24 hours prior to a 6 hour heat (40°C) or cold (25°C) shock. (A-F) Inhibition of farnesylation (FTI or BMS treatment) significantly reduced growth after temperature stress. Growth in GGTI-treated parasites is unchanged after heat or cold shock. (A-F) n = 6; * p≤0.05; ** p≤0.01; *** p≤0.001; **** p≤0.0001, 2-way ANOVA, p-values adjusted for multiple comparisons using Sidak’s multiple comparison test. Abbreviations: control (ctrl), heat shock (hs), cold shock (cs).

**Figure 2.**
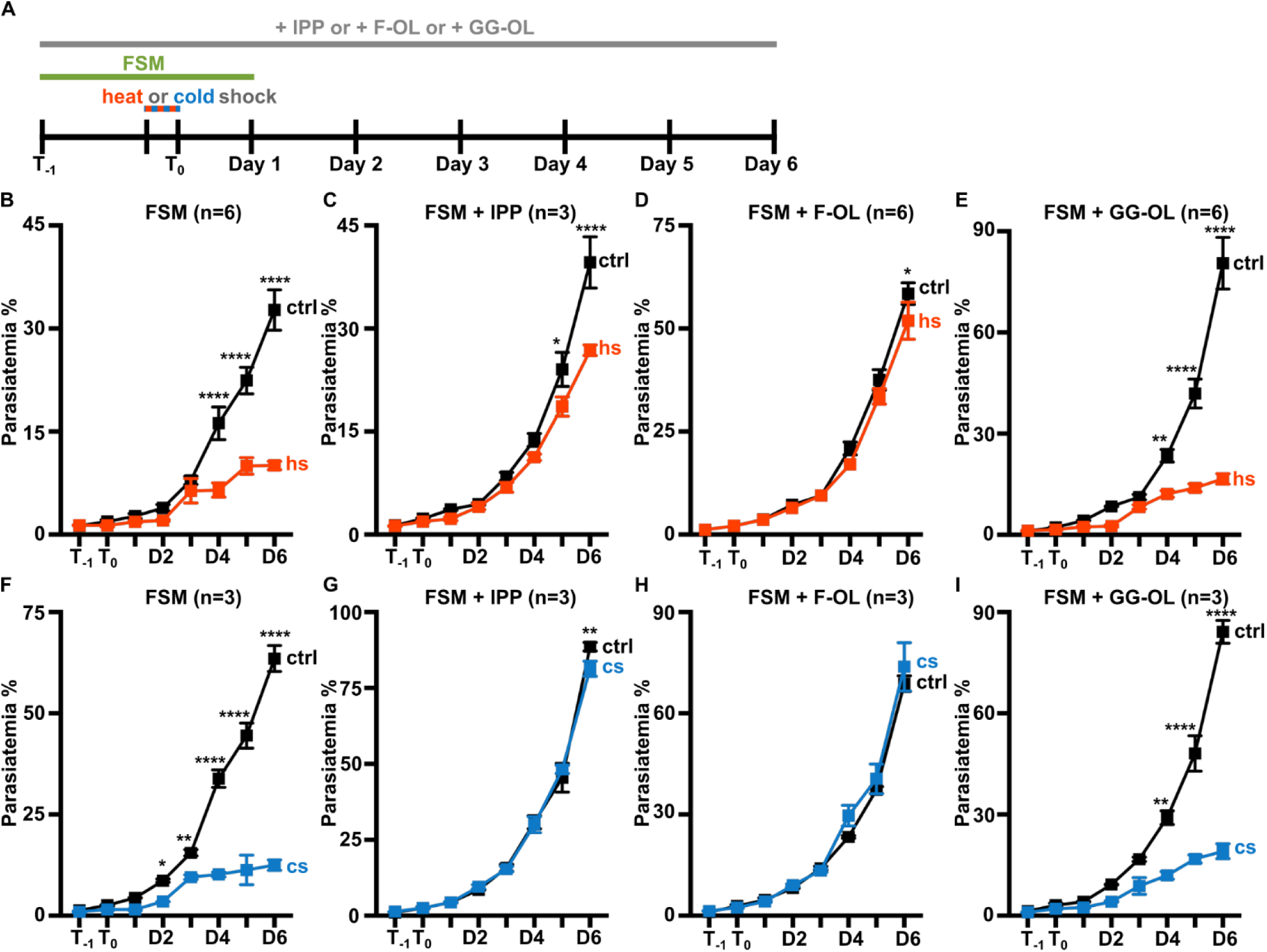
Supplementation with IPP and F-OL rescues growth in FSM-treated parasites after temperature stress. (A) Parasites were treated with FSM (5μM) for 24 hours prior to a 6 hour heat (40°C) or cold (25°C) shock. Cultures were supplemented with isoprenoid products [IPP (250μM), F-OL (10μM), or GG-OL (10μM)] for the entire length of experiment. (B) FSM-treated parasites are sensitive to heat shock. (C,D) Supplementation with IPP (C) or F-OL (D) rescues heat sensitivity. (E) GG-OL supplementation is unable to rescue growth after heat stress. (F-I) IPP or F-OL, but not GG-OL, supplementation is similarly able to rescue FSM-treated parasite growth after cold shock. (B-I) n =3-6; * p≤0.05; ** p≤0.01; *** p≤0.001; **** p≤0.0001, 2-way ANOVA, p-values adjusted for multiple comparisons using Sidak’s multiple comparison test. Abbreviations: control (ctrl), heat shock (hs), cold shock (cs).

### HSP40 is resistant to disruption in P. falciparum

HSP40 is one member of an expanded Hsp40 chaperone family in *P. falciparum*. While the majority of these Hsp40 family members do not have assigned biological functions, HSP40 is considered the canonical member of this family and directly interacts with HSP70 (PF3D7_0818900)^35^. Based on forward genetic screening in *P. falciparum,* HSP40 is predicted to be essential during asexual blood-stage growth^44^. To address the role of HSP40 in asexual development, we sought to disrupt the HSP40 locus directly. Using single-crossover homologous recombination (as previously used to validate the MEP pathway genes, Dxr and IspD)^20, 45^, we successfully integrated a control plasmid into the HSP40 genetic locus (Fig. 3A, B). However, even after 7 months of continuous culture, integration of a disruption construct did not occur (Fig. 3A, C). Our data support an essential role for HSP40 in the parasite during blood-stage growth.

**Figure 3.**
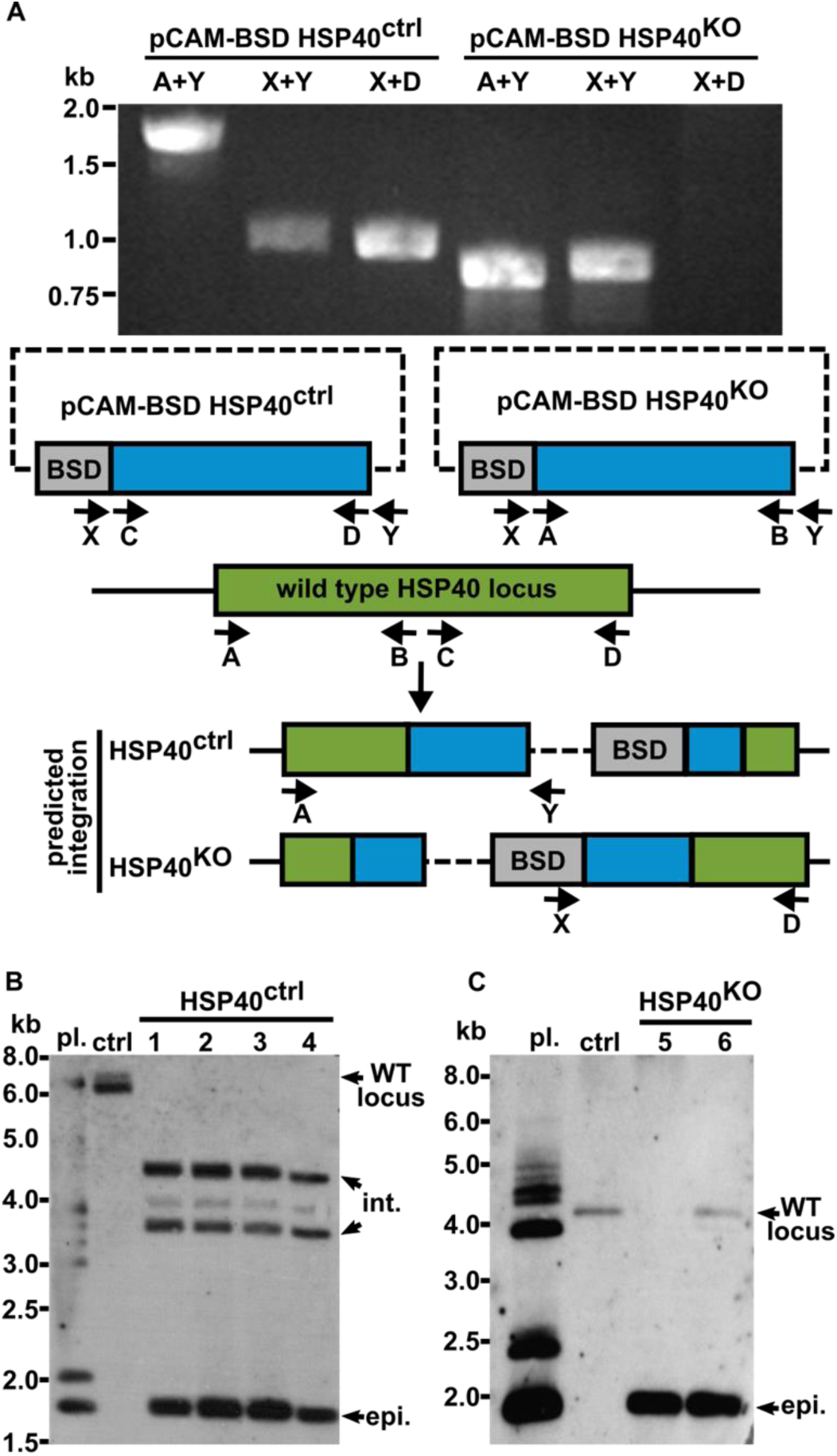
HSP40 locus is not amenable to disruption. Using single-crossover homologous recombination we successfully integrated a control plasmid (pCAM-BSD HSP40^ctrl^) into the HSP40 (PF3D7_1437900) locus. Despite 7 months of continuous culture, integration of a disruption construct (pCAM-BSD HSP40^KO^) was not observed. (A) Genomic DNA isolated from blasticidin-resistant parasites transfected with pCAM-BSD HSP40^ctrl^ or pCAM-BSD HSP40^KO^ was subjected to PCR. Two sets of primers detect episomal plasmid in both cases (ctrl: 1081bp, 1133bp and KO: 902bp, 982bp). Integration events are detected by primers (∼1800bp). Integration of the control plasmid is present, while no integration of the KO plasmid is observed. Representative of three independent transfections. (B,C) Genomic DNA isolated from blasticidin-resistant parasites transfected with pCAM-BSD HSP40^ctrl^ or pCAM-BSD HSP40^KO^ was subjected to Southern blot analysis. (B) Plasmid and genomic DNA was digested with the restriction enzyme SmlI, transferred to membrane, and probed with a 719bp HSP40 control fragment. The control plasmid (pCAM-BSD HSP40^ctrl^) was successfully integrated into the HSP40 locus in four independent transfections (1-4). (C) Plasmid and genomic DNA was digested with the restriction enzyme HpaII, transferred to membrane, and probed with a 693bp HSP40 KO fragment. Integration of a disruption construct (pCAM-BSD HSP40^KO^) was not observed in two independent transfections (5,6) after 7 months of continuous culture.

### HSP40 can stimulate ATPase activity of its co-chaperone HSP70 in vitro

In contrast to the highly expanded HSP40 protein family, the *P. falciparum* genome encodes only 6 Hsp70-like proteins^33, 46^. HSP70 (PF3D7_0818900), the major cytosolic Hsp70 in *P. falciparum,* possesses ATPase activity^47–49^. Select Hsp40 co-chaperones interact with Hsp70 through a protein domain called a J-Domain^50–54^. To determine whether HSP40 interacts with HSP70 to promote ATP hydrolysis, recombinant 6XHis-HSP40 and 6XHis-HSP70 were expressed and purified from *E. coli* (Fig. 4A). We find that the addition of purified recombinant HSP40 stimulates the basal ATP hydrolytic activity of HSP70 (Fig. 4B,C). Increasing the amount of HSP70 in the reaction increases ATP turnover (Fig. 4D); however, the addition of HSP40 increases the ATPase activity of HSP70 nearly 3-fold (Fig. 4C). These data, along with previous observations by Botha et al. (2011), provide evidence of a functional interaction between HSP40 and HSP70.

**Figure 4.**
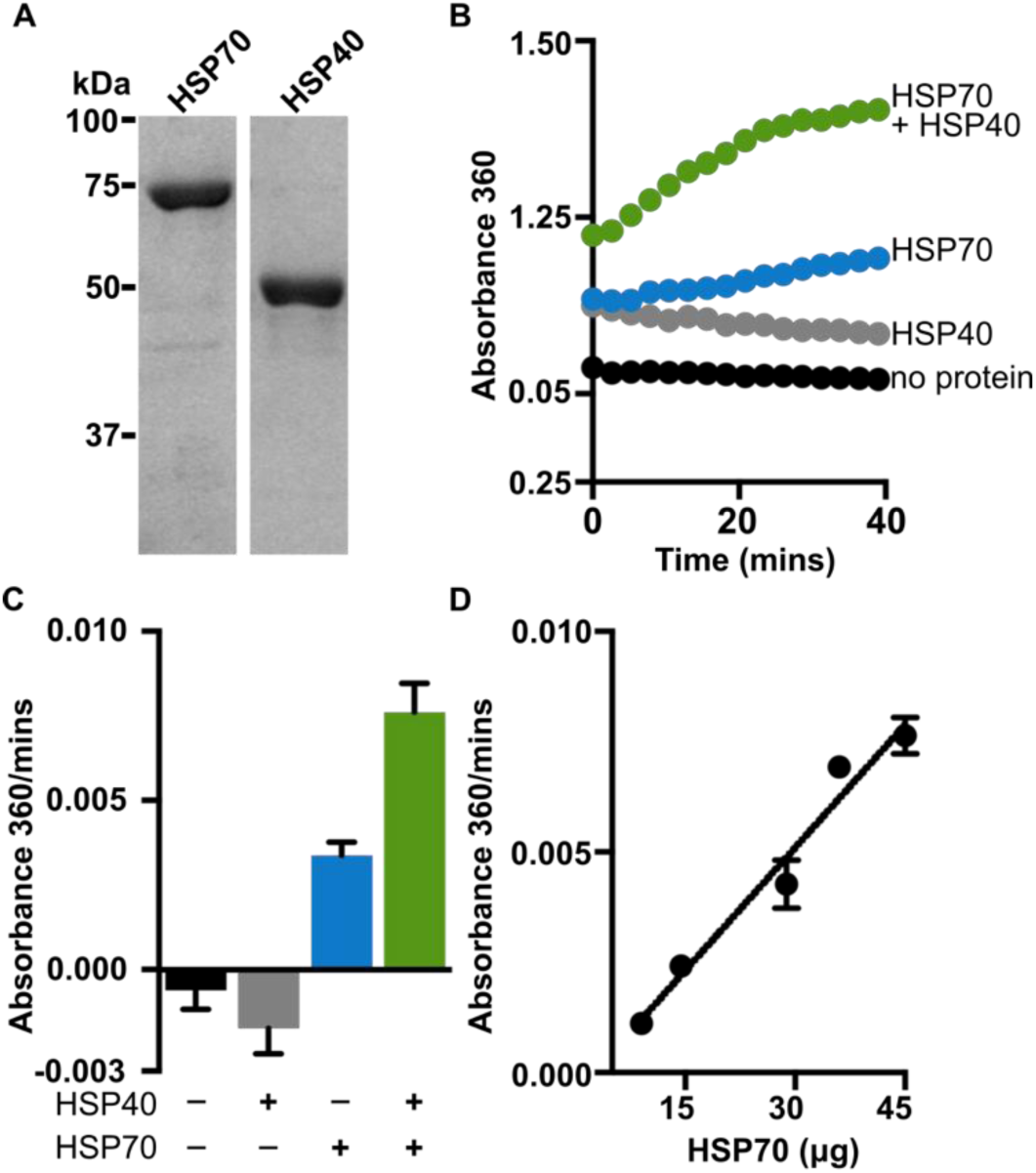
ATPase activity of purified HSP70. (A) Recombinant HSP70 (73 kDa) and HSP40 (48 kDa) were purified from *E. coli.* Proteins are visualized on SDS gel with Coomassie blue stain (B) The ATPase activities of 45 μg of HSP70 was measured every 12 seconds for 40 minutes. HSP70 had increased ATPase activity in the presence of HSP40 (8.9 μg; green) than alone (blue). HSP40 (gray) and no protein (black) were used as controls, n=4. (C) The slopes of the lines in (B). (D) The rate of ATP turnover increases linearly with increased HSP70 concentration, n=2.

### Robust HSP40 membrane association requires IPP synthesis and protein farnesylation

Using purified recombinant 6XHis-HSP40 (Fig. 4A), polyclonal antisera were generated. Immunoblotting with anti-HSP40 antisera reveals a single band in parasite lysate, which is not present in uninfected RBCs (Fig. S3A,B). Antisera specificity was confirmed by immunoprecipitation (IP) and mass spectrometry, demonstrating exclusive immunoprecipitation of HSP40 without cross-reactivity to other Hsp40 family members in *P. falciparum* (Fig. S3C).

Since heat shock proteins help mediate export of parasite proteins through the PTEX complex^55–57^, we performed immunofluorescence localization of HSP40 in asexual *P. falciparum*. HSP40 localizes to the parasite cytosol in trophozoite-stage parasites and is not detected in the host cell (Fig. 5A). Immuno-electron microscopy (immuno-EM) was performed to further characterize the subcellular localization of HSP40. We find that HSP40 is distributed throughout the cytosol and excluded from the nucleus and host cell cytoplasm (Fig. 5B). Because farnesylation marks small GTPases for localization to the cytosolic face of the endoplasmic reticulum (ER)^58, 59^, we predicted that some portion of HSP40 may be ER-localized. In addition to anti-HSP40, parasites were labeled with anti-PDI, an established ER marker^60, 61^. A subset of HSP40 is localized alongside PDI in the ER (Fig. 5B). These data indicate that HSP40 is both cytosolic and ER-localized.

HSP40 is found in both the soluble (data not shown) and membrane fractions (Fig. 6A,B), following detergent fractionation of parasite lysate. While overall levels of HSP40 protein remain unchanged with inhibition of IPP synthesis or farnesylation, the percentage of membrane-associated HSP40 is significantly reduced (Fig. 6C,D). Treatment with farnesylation inhibitors (such as FTI or BMS), but not the geranylgeranylation inhibitor GGTI, reduces membrane-associated HSP40 (Fig S4). Subcellular localization of HSP40 is changed after treatment with either FSM or FTI (Fig. 6E,F). Quantification of immuno-EM micrographs confirms a reduced membrane association of HSP40 in inhibitor-treated cells (Fig. 6G,H). Overall, these observations provide evidence that a subset of HSP40 is membrane-associated and that this membrane association requires isoprenoid synthesis and farnesylation.

**Supplemental Figure 3.**
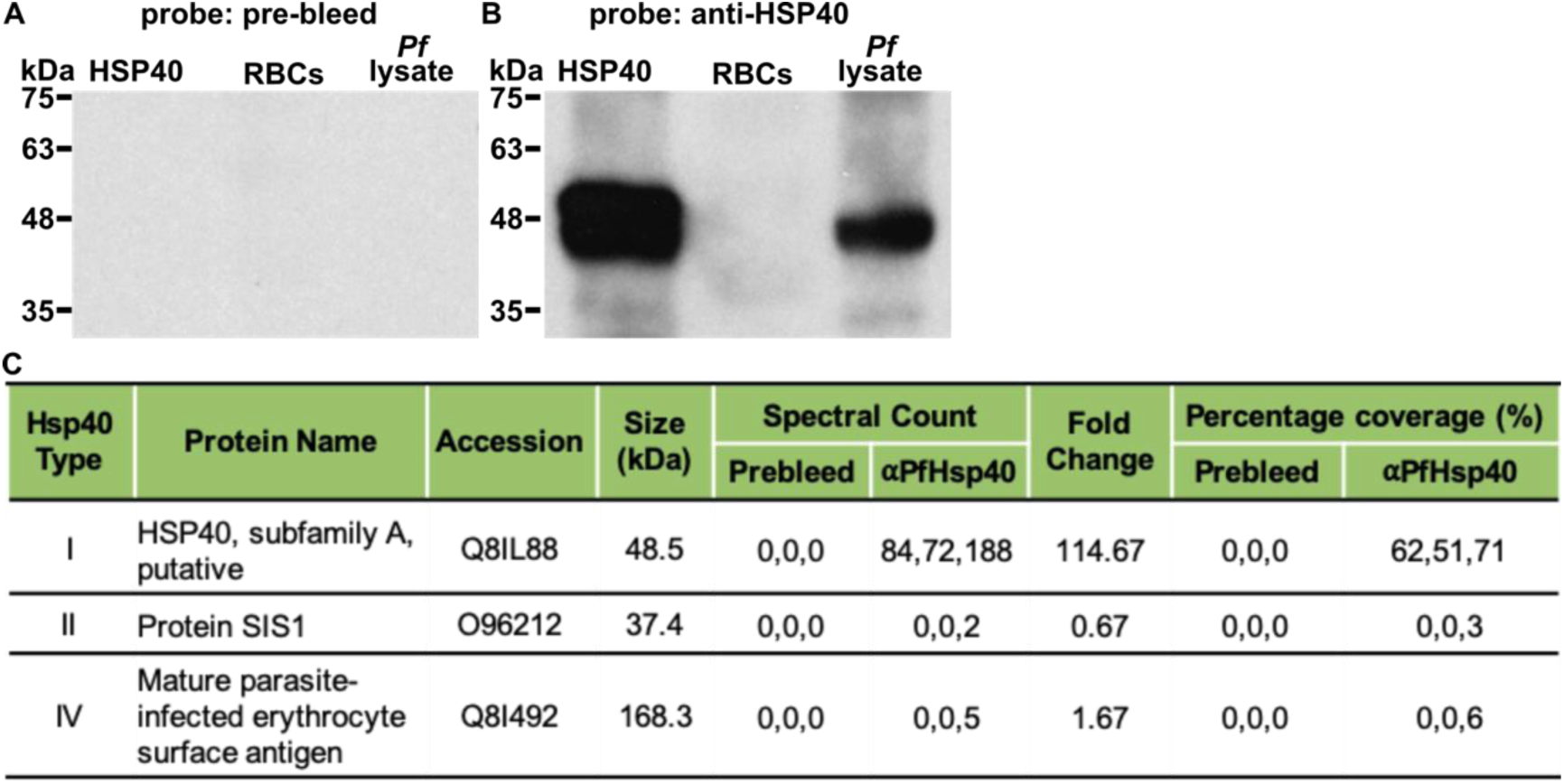
Polyclonal anti-HSP40 antisera is specific for HSP40. (A) Pre-bleed antisera immunoblot of recombinant protein, RBC lysate, and wildtype *P. falciparum* lysate. (B) Anti-HSP40 immunoblot of recombinant protein, RBC lysate, and wildtype *P. falciparum* lysate. A single band is observed at 48 kDa in parasite lysate corresponding to the expected size of HSP40. (A,B) Pre-bleed and Anti-HSP40 were used at 1:5,000. (C) Spectral counts from Hsp40s identified by mass spectrometry after IP of parasite lysate with pre-bleed antisera or anti-HSP40. Fold change is the ratio of the average spectral counts between anti-HSP40 and pre-bleed. HSP40 is the only Hsp40 protein with a robust fold change and protein coverage.

**Figure 5.**
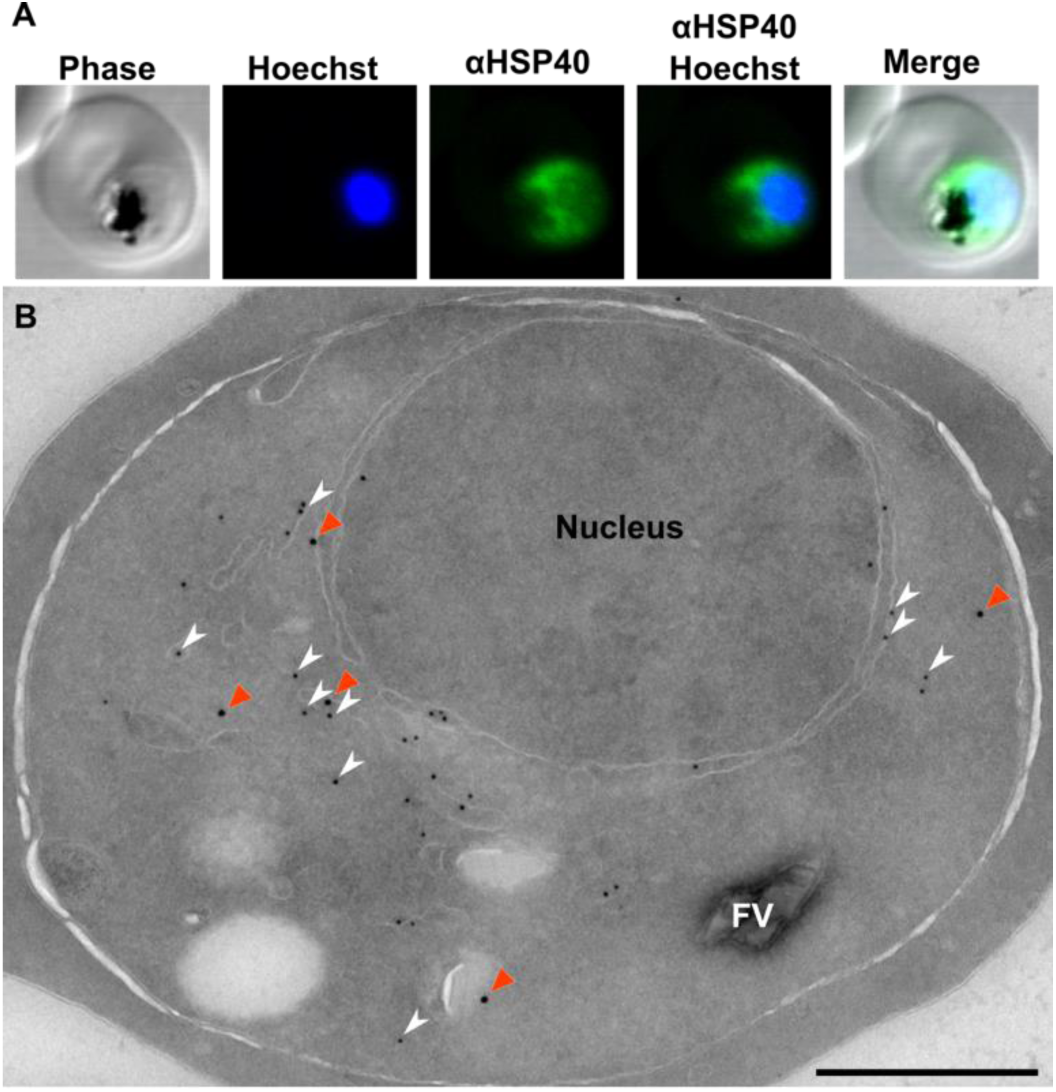
Localization of HSP40 in *P. falciparum.* (A) Immunofluorescence confocal microscopy of trophozoite, stained with anti-HSP40 (1:5,000) and Hoechst 33258 nuclear stain. HSP40 appears cytosolic. (B) Electron micrograph of Immunolabeling: primary, rabbit anti-HSP40 (1:250), mouse anti-PDI (1:100); secondary, goat anti-rabbit IgG 18nm colloidal gold, goat anti-mouse 12nm. HSP40 (orange arrowheads) looks cytosolic in the parasites, with some apparent membrane association. A portion of HSP40 co-localizes with PDI, an established ER marker (white arrowheads). Scale, 500 nm.

**Figure 6.**
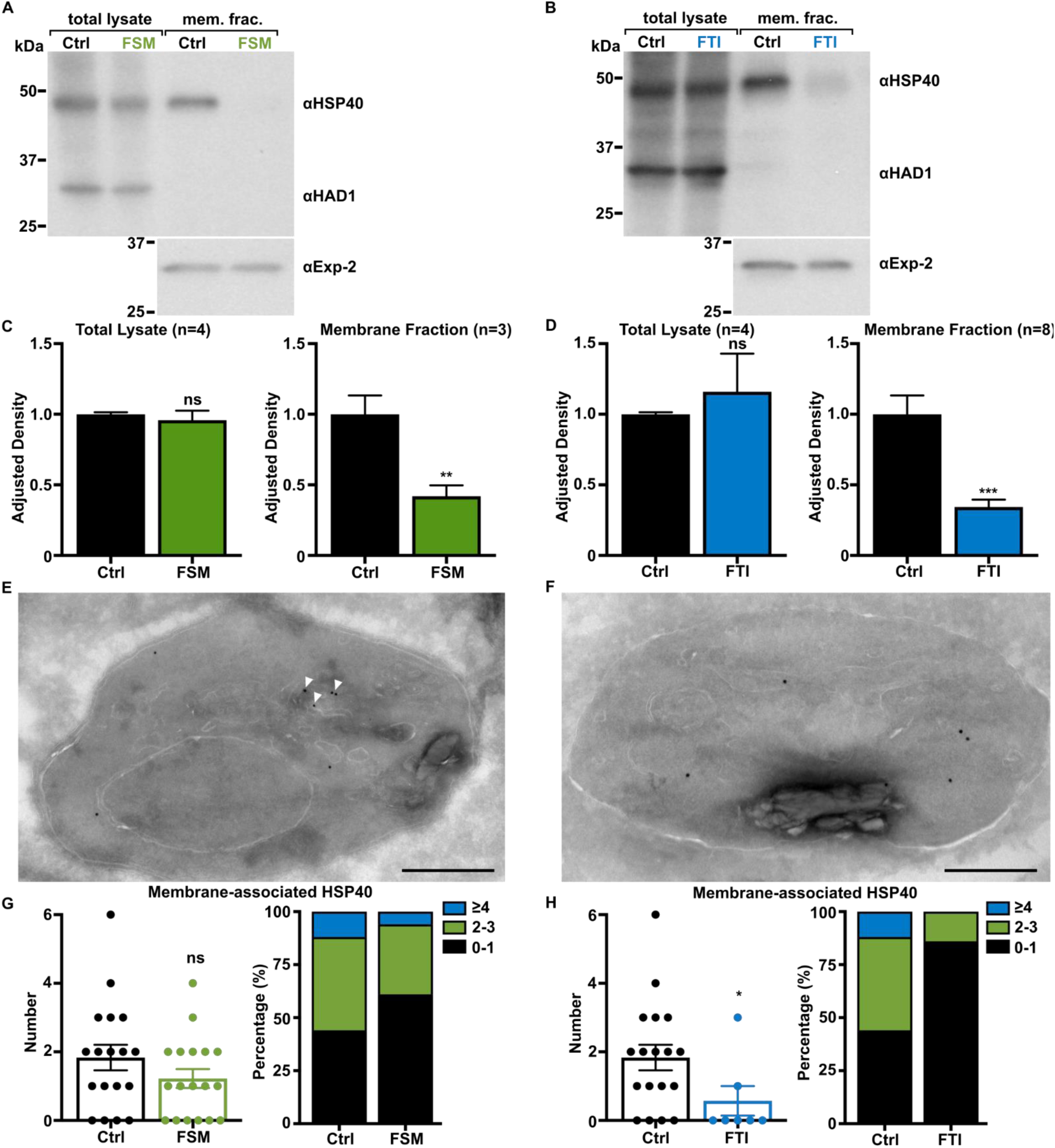
Inhibition of either IPP synthesis or protein farnesylation results in reduced membrane-association of HSP40. (A,B) Representative anti-HSP40 immunoblots of control and FSM (20μM) treated (A) or FTI (10μM) treated (B) *P. falciparum* total lysate and membrane fractions. (C,D) Quantification of several immunoblots adjusted to loading control. HSP40 is significantly reduced in the membrane fraction after inhibition of IPP synthesis (C) and inhibition of farnesylation (D). Anti-HAD1 and Anti-Exp-2, loading controls for total lysate and membrane fractions, respectively. ** p≤0.01, *** p≤0.001 unpaired t-test with Welch’s correction. (E-H) HSP40 membrane association is reduced after FSM and FTI treatment. Apparent membrane-associated HSP40 (10nm gold particles, arrowheads) is reduced after inhibition of IPP synthesis and farnesylation. The number of membrane-associated HSP40 per micrograph is quantified for control and treated parasites (G,H). A single control cohort was quantified. A decrease in the number of membrane-associated HSP40 particles is observed. * p≤0.05, unpaired t-test with Welch’s correction. Scale, 500 nm.

**Supplemental Figure 4.**
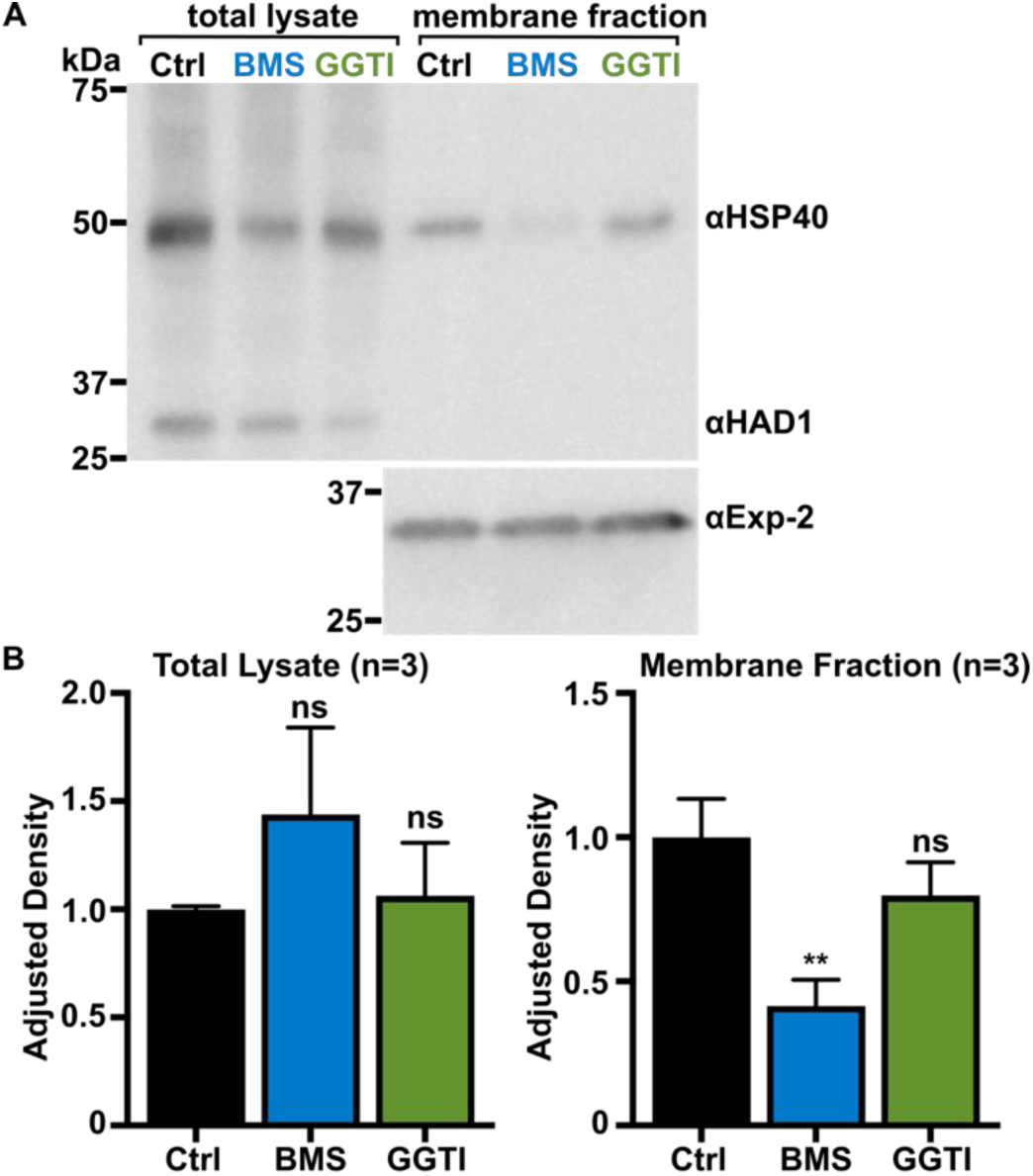
Farnesylation, not geranylgeranylation, mediates membrane association of HSP40. (A) Representative anti-HSP40 immunoblots of control, BMS (200nM) and GGTI (2μM) treated *P. falciparum* total lysate and membrane fractions. (B) Quantification of several immunoblots adjusted with loading control. HSP40 is significantly reduced in the membrane fraction after inhibition of farnesylation (BMS) and not geranylgeranylation (GGTI). Anti-HAD1 and Anti-Exp-2, loading controls for total lysate and membrane fractions, respectively. ** p≤0.01, unpaired t-test with Welch’s correction.

### Palmitoylation contributes to HSP40 membrane association, but not thermotolerance

Farnesylation is not the only post-translational modification that is expected to bring HSP40 to the membrane, as HSP40 is also palmitoylated^62^. To evaluate the role of palmitoylation in the membrane association of HSP40 and parasite thermotolerance, we employed the palmitolyation inhibitor, 2-bromopalmitate (2BP)^63^. While overall levels of HSP40 remain unchanged upon inhibition of palmitoylation, the proportion of membrane-associated HSP40 is significantly reduced (Fig. 7A,B). Combined treatment with 2BP and FTI (to inhibit both palmitoylation and farnesylation) further reduces the levels of membrane-associated HSP40, compared to either treatment alone (Fig. 7A,B). We tested whether inhibition of palmitoylation influenced parasite growth following heat (40°C) or cold (25°C) stress. Consistent with our previous observations, untreated parasites readily recover after modest heat or cold shock (Fig. 7C, G), while inhibition of farnesylation significantly impairs recovery after temperature stress (Fig. 7D,H). In contrast, treatment with 2BP does not sensitize parasites to temperature stress, as growth is unchanged following heat and cold shock (Fig. 7E,I). Loss of protein palmitoylation was also not protective, as parasites treated with both FTI and 2BP remained sensitive to temperature shock (Fig. 7F,J). Therefore, farnesylation, but not palmitoylation, is required for parasite thermotolerance. Overall, these data indicate that protein prenylation plays a critical role in parasite thermotolerance and is necessary, but not sufficient, for membrane-association of HSP40.

**Figure 7.**
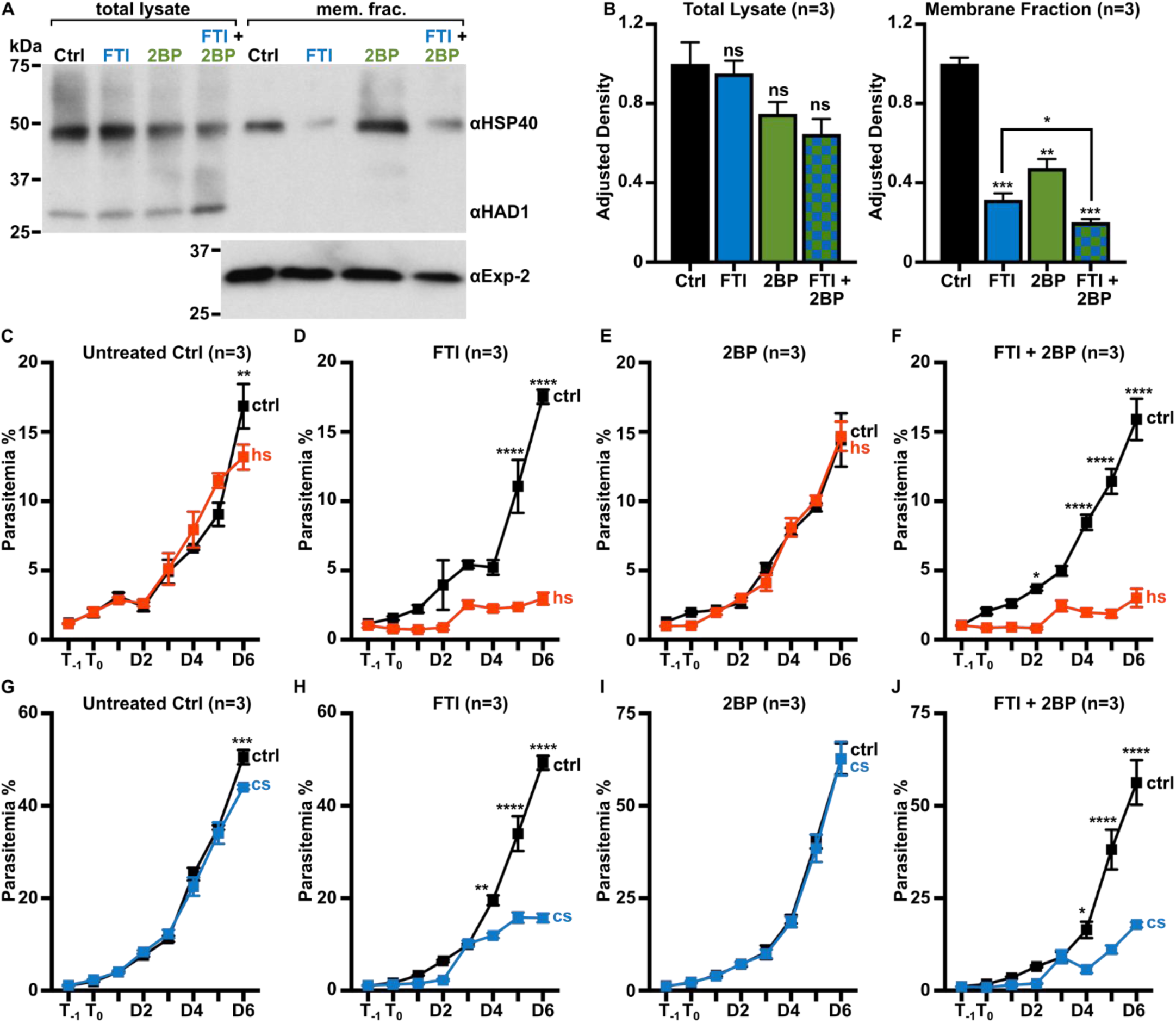
Both farnesylation and palmitoylation contribute to HSP40 membrane-association, but only farnesylation is required for thermotolerance. (A) Representative anti-HSP40 immunoblots of control, FTI (10μM), 2BP (100μM), and combination of FTI- and 2BP-treated *P. falciparum* total lysate and membrane fractions. (B) Quantification of several immunoblots adjusted with loading control. The membrane-associated proportion of HSP40 is significantly reduced upon inhibition of farnesylation (FTI) or palmitoylation (2BP). Inhibition of both farnesylation and palmitoylation (FTI + 2BP) further reduces HSP40 membrane-association, as compared to single inhibitor treatment. Anti-HAD1 and Anti-Exp-2, loading controls for total lysate and membrane fractions, respectively. n = 3; * p≤0.05; ** p≤0.01; *** p≤0.001 unpaired t-test with Welch’s correction. (C-J) Parasites were treated with either FTI (10μM), 2BP (100μM), or both prior to heat (40°C) or cold (25°C) shock. FTI-treated parasite growth is significantly reduced after heat (D) and cold shock (H). Growth in 2BP-treated parasites is unchanged after heat or cold shock. (E,I). Parasites treated with both FTI and 2BP were sensitive to temperature stress (F,J). n = 3; * p≤0.05, ** p≤0.01; *** p≤0.001; **** p≤0.0001, 2-way ANOVA, p-values adjusted for multiple comparisons using Sidak’s multiple comparison test. Abbreviations: control (ctrl), heat shock (hs), cold shock (cs).

### HSP40 interacts with GAPDH and components of the cytoskeleton in an IPP-dependent manner

HSP40 is presumed to interact with numerous co-chaperones and client proteins. Protein prenylation is known to drive association with the ER and is likely to alter accessibility to client proteins^64^. To understand the mechanism by which loss of protein prenylation might impact HSP40 function and parasite thermotolerance and survival, we sought to identify HSP40-interacting proteins in the presence and absence of protein prenylation. We used immunoprecipitation and mass spectrometry to identify HSP40-interacting proteins from parasites under normal prenylation conditions (wild type controls) and when prenylation is impaired (FSM or FTI treatment).

We find that HSP40-interacting proteins mediate a number of essential biological functions in the parasite, including cytoskeleton organization, glycolysis, translation, and pathogenesis (Table S1). HSP40 protein interactors are significantly enriched in the following biological processes: translation, glycolytic process, and gluconeogenesis (Fig. 8A). Interestingly, inhibition of farnesylation does not prevent the enrichment of translation protein interactors (Fig. 8A). The interaction of HSP40 with cytoskeleton organizational components (Tubulin ⍺ chain, Tubulin β chain, and Actin-1) is significantly reduced upon inhibition of IPP synthesis and farnesylation (Fig. 8B,C). Similarly, the interaction of HSP40 with the glycolytic enzyme GAPDH is significantly reduced after treatment with either FSM or FTI (Fig. 8B,C). The interaction between HSP40 and its co-chaperone HSP70 is significantly increased after inhibition of farnesylation, but not IPP synthesis. Therefore, the interaction between HSP40 and its co-chaperone HSP70 may be independent of farnesylation, as HSP70 appears to interact with the unfarnesylated form of HSP40.

**Figure 8.**
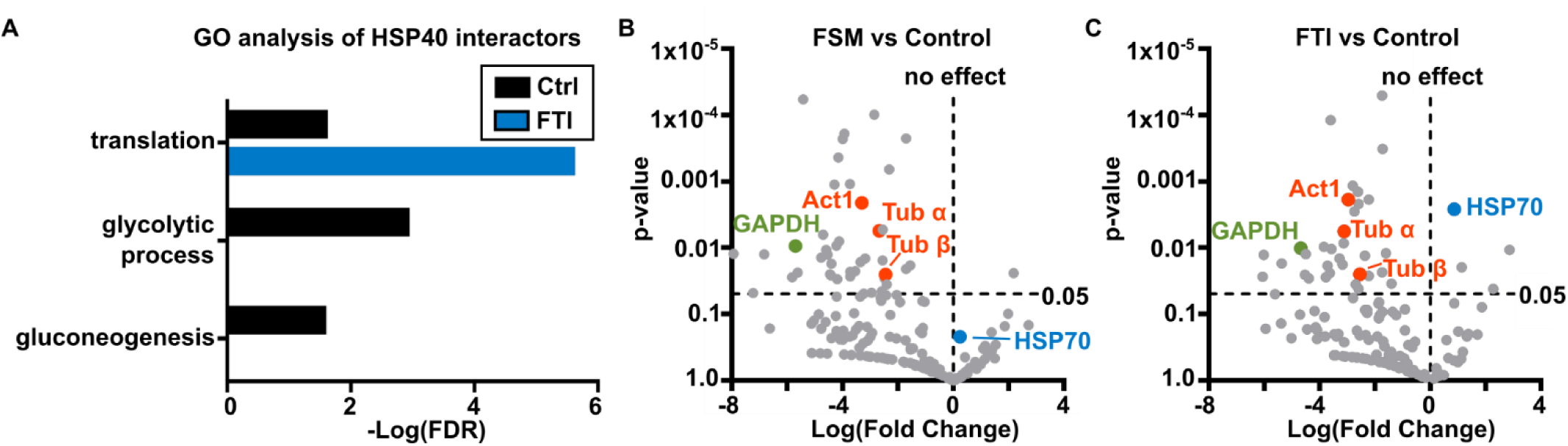
Both IPP synthesis and protein farnesylation influence interactions of HSP40 with GAPDH and cytoskeleton components. (A) Gene ontology analysis reveals three significant HSP40-associated biological processes in asexual *P. falciparum.* All significant associations are lost in FSM-treated parasites. (B,C) The interactions of HSP40 with Tubulin α chain [Tub α; Q6ZLZ9], Tubulin chain [Tub β; Q7KQL5], Actin-1 [Act1; Q8I4X0], and GAPDH [Q8IKK7] are significantly reduced after inhibition of IPP synthesis and/or farnesylation. Interestingly, interactions with HSP70 [Q8IB24] are significantly increased after farnesylation inhibition, but not after inhibition of IPP synthesis. Protein-protein interactions were determined by mass spectrometry after IP of parasite lysate with anti-HSP40. Results for FSM (5μM)- and FTI (10μM)-treated parasites are compared to untreated controls. n = 3; multiple unpaired t-tests.

**Supplemental Table 1.** HSP40 interactors during the intraerythrocytic *P. falciparum*.

| Untreated Control |  |  |  |  |  | FSM |  | FTI-277 |  |
| --- | --- | --- | --- | --- | --- | --- | --- | --- | --- |
| Gene Ontology - Biological Process | Protein Name | Accession Number | Molecular Weight (kDa) | Average Peptide Count | Average Sequence Coverage (%) | fold change | p-value | fold change | p-value |
| chromosome organization | Histone H2B | Q8IIV1 | 13 | 11.0 | 32.0 | 0.053 | 0.0093 | 0.070 | 0.0097 |
| cytoskeleton organization | Tubulin beta chain | TBB | 50 | 21.7 | 36.0 | 0.184 | 0.0253 | 0.171 | 0.0250 |
|  | Tubulin alpha chain | Q6ZLZ9 | 50 | 17.3 | 32.7 | 0.157 | 0.0056 | 0.116 | 0.0056 |
|  | Actin-1 | ACT1 | 42 | 12.3 | 20.0 | 0.101 | 0.0021 | 0.128 | 0.0019 |
| Glycolytic process | Glyceraldehyde-3-phosphate dehydrogenase | Q8IKK7 | 37 | 30.0 | 49.0 | 0.019 | 0.0094 | 0.039 | 0.0101 |
|  | Phosphoglycerate kinase | PGK | 45 | 17.0 | 21.7 | 0.053 | ns | 0.016 | ns |
|  | Enolase | ENO | 49 | 15.3 | 25.0 | 0.047 | ns | 0.059 | ns |
|  | Pyruvate kinase | C6KTA4 | 56 | 13.3 | 19.1 | 0.053 | ns | 0.071 | ns |
| nucleocytoplasmic transport | GTP-binding nuclear protein | Q7KQK6 | 25 | 10.3 | 36.0 | 0.488 | 0.0655 | 0.529 | 0.0845 |
| ornithine metabolic process | Ornithine aminotransferase | OAT | 46 | 13.3 | 14.7 | 0.029 | ns | 0.024 | ns |
| pathogenesis | Merozoite surface protein 1 | Q8I0U8 | 196 | 13.0 | 7.5 | 0.043 | 0.0126 | 0.044 | 0.0125 |
| phosphatidylcholine biosynthetic process | Phosphoethanolamine N-methyltransferase | Q8IDQ9 | 31 | 12.0 | 34.7 | 0.007 | 0.0481 | 0.020 | 0.0501 |
| Response to heat and unfolded protein | Heat shock protein 70 | Q8I2X4 | 72 | 43.7 | 20.0 | 0.435 | ns | 0.577 | ns |
|  | Heat shock protein 70 | Q8IB24 | 74 | 36.0 | 44.7 | 1.184 | ns | 1.822 | 0.0026 |
| Translation | Elongation factor 1-alpha | Q8I0P6 | 49 | 40.7 | 43.7 | 0.306 | 0.0002 | 0.303 | 0.0003 |
|  | 40S ribosomal protein S19 | Q8IFP2 | 20 | 30.0 | 48.0 | 0.263 | 0.0656 | 0.273 | 0.0691 |
|  | 40S ribosomal protein S18 | Q8IIA2 | 18 | 14.3 | 45.3 | 0.083 | 0.0969 | 0.055 | 0.0898 |
|  | 40S ribosomal protein S9 | Q8I3R0 | 22 | 12.0 | 23.3 | 0.138 | 0.0001 | 0.299 | 0.0001 |
|  | 40S ribosomal protein S27 | Q8IEN2 | 9 | 11.0 | 35.0 | 0.047 | 0.0174 | 0.076 | 0.0186 |
|  | 40S ribosomal protein S10 | Q8IBQ5 | 16 | 10.3 | 21.3 | 0.004 | 0.0122 | 0.015 | 0.0127 |

### HSP40 farnesylation influences GAPDH localization but not its glycolytic function

The glycolytic enzyme GAPDH (PF3D7_1462800) has a well-known crucial function in the glycolytic breakdown of glucose to produce ATP in the parasite. In other systems, GAPDH also has an important non-catalytic role in mediating ER-to-Golgi endomembrane trafficking through interaction with microtubules^65–67^. Using purified recombinant 6XHis-GAPDH, specific polyclonal antisera were generated (Fig. S5A,B). As observed for HSP40, we find that GAPDH is, in part, membrane-associated and this membrane association is also dependent on IPP synthesis and farnesylation (Fig. 9A-C).

To understand whether interrupting IPP synthesis and protein prenylation directly impacts GAPDH function, we quantified glycolytic intermediates in the presence and absence of IPP synthesis and protein prenylation. We find that the cellular levels of the GAPDH substrate [glyceraldehyde 3-phosphate (GAPD)] and its immediate downstream product [2/3-phosphoglycerate (2/3-PGA)], are unchanged after treatment with either FSM or FTI (Fig 9D-F). Substrate availability to the pentose phosphate pathway also does not change upon FSM or FTI treatment (Fig S6). Our data indicate that farnesyl-HSP40 does not directly influence the catalytic function of GAPDH in glycolysis. Our findings instead suggest that the prenylation-dependent interaction between HSP40 and GAPDH on the endoplasmic reticulum may mediate non-catalytic (so-called “moonlighting”) biological roles of GAPDH, such as its known role in endomembrane trafficking^65, 66, 68–76^.

**Figure 9.**
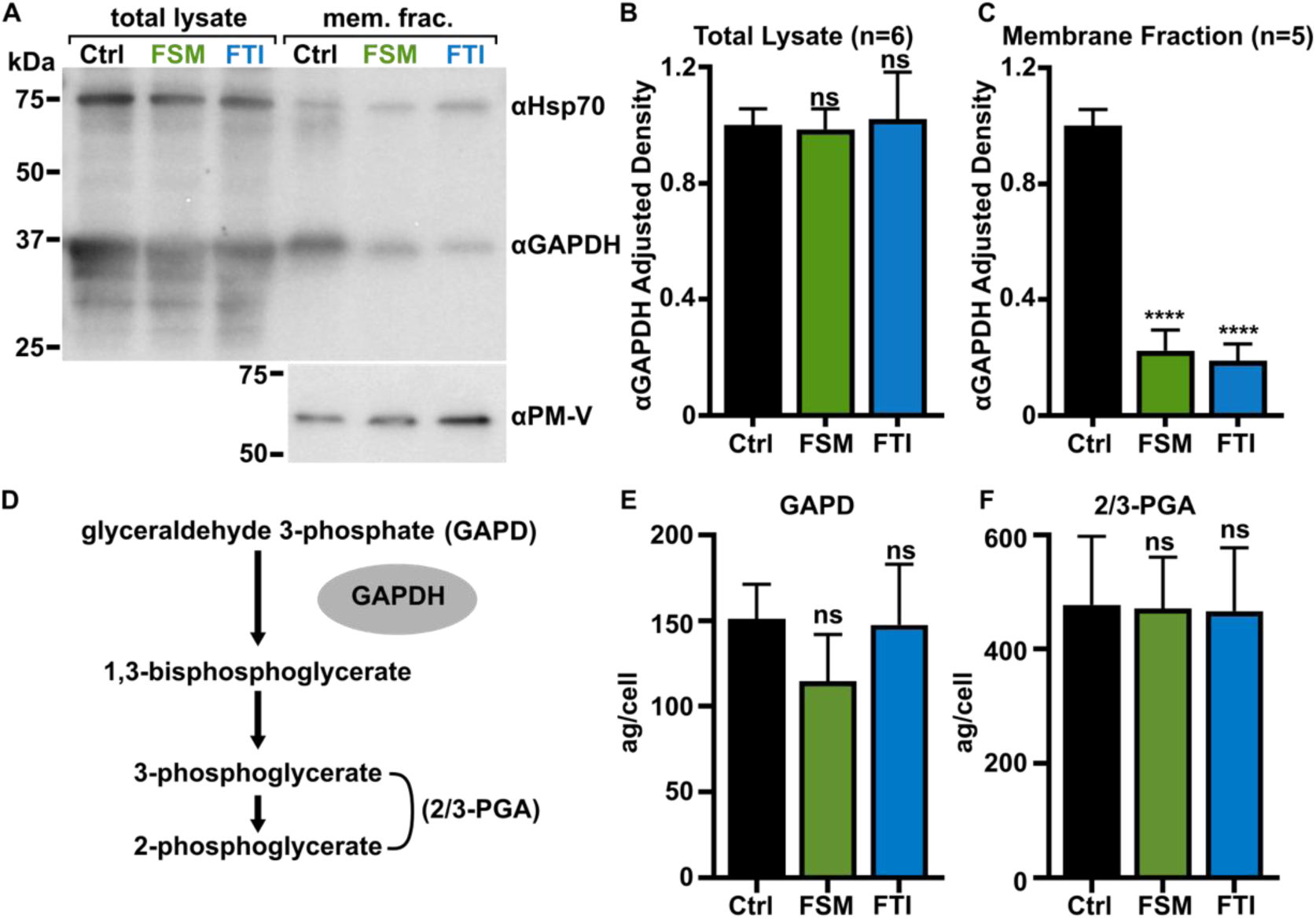
Localization of GAPDH, but not its glycolytic function, are IPP- or farnesylation-dependent. (A-C) Inhibition of IPP synthesis and farnesylation reduced membrane association of GAPDH. (A) Representative anti-GAPDH immunoblots of control, FSM (20μM) and FTI (10μM) treated *P. falciparum*. (B,C) Quantification of several immunoblots adjusted with loading control. Anti-Hsp70 (1:5,000) and anti-PM-V (1:500) were used as loading controls for total lysate and membrane fractions, respectively. n = 5-6; **** p≤0.0001, unpaired t-test with Welch’s correction. (D) GAPDH catalyzes the conversion of glyceraldehyde 3-phosphate (GAPD) to 1,3-bisphosphoglycerate, which is further reduced to 2/3-phosphoglycerate (2/3-PGA). (E,F) Levels of GAPD and 2/3-PGA were measured by liquid chromatography with tandem mass spectrometry and normalized based on parasitemia of each individual sample to give concentration per cell. The amount of GAPD and 2/3-PGA is unchanged after treatment with FSM (5μM) or FTI (10μM). n=3, unpaired t-test with Welch’s correction.

**Supplemental Figure 5.**
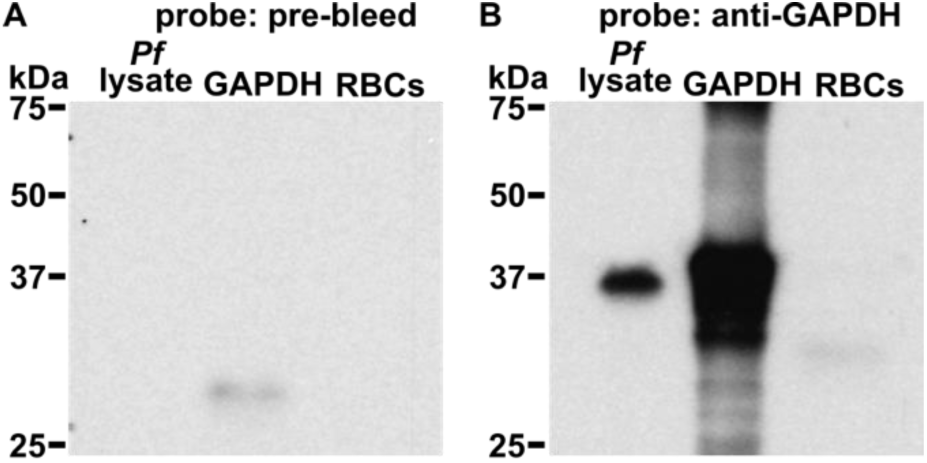
Polyclonal anti-GAPDH antisera. (A) Pre-bleed antisera immunoblot of wildtype *P. falciparum* lysate, recombinant protein, and RBC lysate. (B) Anti-GAPDH immunoblot of wildtype *P. falciparum* lysate, recombinant protein, and RBC lysate. A single band is observed at 37 kDa in parasite lysate corresponding to the expected size of GAPDH. (A,B) Pre-bleed and Anti-GAPDH were used at 1:1,000.

**Supplemental Figure 6.**
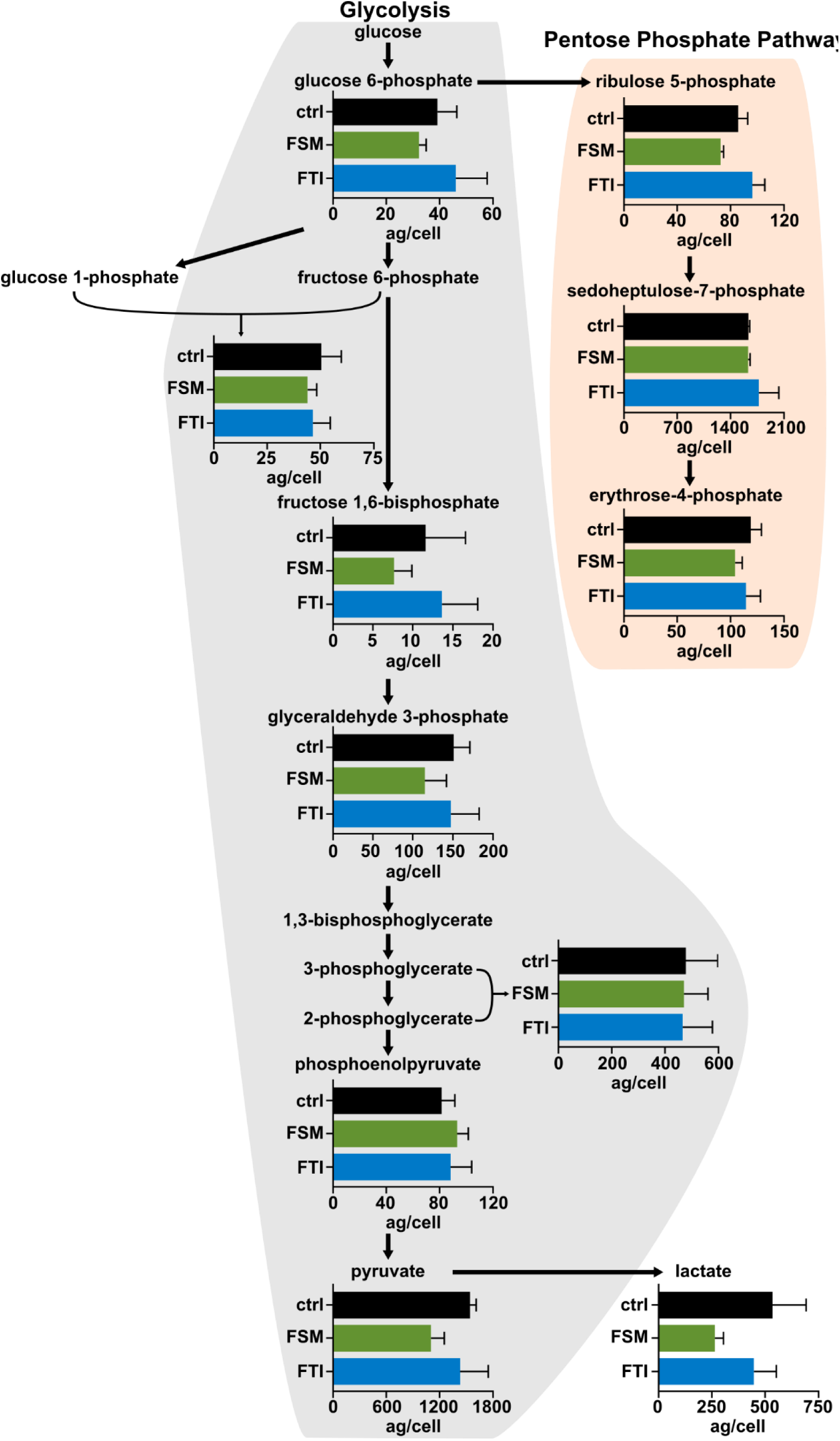
Glycolytic and pentose phosphate pathway metabolite levels remain constant under IPP and farnesylation-deficient conditions. Levels of glycolytic and pentose phosphate pathway intermediates were measured by liquid chromatography with tandem mass spectrometry and normalized based on parasitemia of each individual sample to give concentration per cell. No significant changes are observed after treatment with FSM (5µM) or FTI (10µM). n=3, unpaired t-test with Welch’s correction.

## Discussion

Temperature change is a critical environmental signal and an integral part of the *P. falciparum* life cycle. In addition, antimalarial activity of the first-line artemisinin-based therapies may be sensitive to heat or cold stress ^6, 8^. The mechanisms by which the parasite copes with thermal stress are not well understood, and the inherent host temperature fluctuations during clinical malaria may be exploited to improve malaria treatment. In this study, we establish that survival during heat or cold shock in *P. falciparum* requires both *de novo* isoprenoid biosynthesis and the post-translational 15-carbon isoprenyl modification called farnesylation. Chemical inhibitors that reduce protein farnesylation—either by reducing isoprenoid biosynthesis or farnesyltransferase activity—sensitize *P. falciparum* to otherwise non-lethal temperature stresses. Such inhibitors might be particularly valuable in the setting of malarial fever, a nearly universal characteristic of symptomatic *P. falciparum* infection^77^.

Housed within the apicoplast organelle, isoprenoid biosynthesis through the MEP pathway is necessary for intraerythrocytic development of *P. falciparum*^18–20^. Isoprenyl modification of proteins is a key essential function of the MEP pathway, as prenylation itself is essential for asexual development^23–27^. However, the pleiotropic downstream effects brought on by loss of protein prenylation have not been fully elucidated. Kennedy et al. recently proposed a mechanistic model of “delayed death,” a phenotype of parasite demise during the second erythrocytic cycle following drug treatment, exhibited by antimalarials that target apicoplast maintenance. In this model, disruption of Rab protein geranylgeranylation and interruption of subsequent cellular trafficking is responsible for the growth arrest caused by loss of IPP production or protein prenylation^78^. Geranylgeranylated Rab GTPase family members comprise the majority of prenylated proteins in *P. falciparum*^30, 31^. However, our data suggest that the phenotype in malaria parasites caused by loss of isoprenoid biosynthesis or protein prenylation is more complicated and interruption of protein farnesylation (even when geranylgeranylation is preserved) also plays an important role. Both geranylgeranylation of Rab GTPases and farnesylation – likely of HSP40 – are key to the essential nature of prenylation in *P. falciparum*.

Our data also suggest the presence of several distinct cellular pools of HSP40 that have different post-translational modifications, subcellular localizations, and client protein interactomes. Maximal membrane association of HSP40 requires post-translational palmitoylation, but palmitoylation alone is insufficient for the biological function of HSP40 in thermotolerance. Thus, palmitoylation may guide HSP40 to a membrane compartment that is different than that of farnesyl-HSP40. Similar findings have been described for mammalian Ras proteins, which, as for HSP40, are both farnesylated and palmitoylated. For Ras proteins, both post-translational modifications are needed for stable plasma membrane association^79^. While our data indicate that the farnesylated pool of HSP40 functions in thermotolerance, pamitoyl-HSP40 appears to have other, yet unidentified, functions in malaria parasites.

While heat shock proteins are found across taxa, not all organisms possess prenylated Hsp40s^80^. HSP40 orthologs in human, yeast, *Plasmodium* spp. (*P. vivax*, *P. yoelii, P. chabaudi,* and *P. berghei*), and plants (*Arabidopsis thaliana* and *Atriplex nummularia*) have been experimentally demonstrated to be prenylated or are predicted to be prenylated based on the presence of canonical C-terminal prenylation sequence motifs^36, 81–83^. It is unclear how the function of Hsp40 orthologs might differ in organisms that lack prenylated Hsp40s. However, our data, in conjunction with studies in yeast and plants, provide strong evidence that prenylation of Hsp40 controls critical functions in the cell including growth after temperature or drought stress, localization within the cell, and interactions with client proteins and signaling pathways ^36–38, 82, 84, 85^. Together, these observations suggest that prenylation of Hsp40 is a marker that distinguishes a distinct functional subclass of Hsp40s, which appear to play similar cellular roles in animals, plants, and fungi, and have essential biological functions that are relatively conserved across the domains of life.

HSP40 is one of only 4 farnesylated proteins in *P. falciparum* and the sole farnesylated heat shock protein^30, 31^. We find that farnesylation controls membrane association of HSP40 on the endoplasmic reticulum and modulates access to its client proteins, such as GAPDH and tubulin subunits, which are required for membrane trafficking in the early secretory pathway. In other systems, GAPDH modulates Rab-dependent vesicular transport and binds directly with microtubule components including tubulin α, in a role that does not rely on its enzymatic activity^65, 66, 68, 69, 71–76^. In malaria parasites, vesicular transport is dramatically disrupted with inhibition of IPP synthesis or protein prenylation ^23, 78^. Therefore, we propose a novel cellular model whereby farnesyl-HSP40 functions in vesicular trafficking, through its interactions with both GAPDH and the cytoskeleton (Fig. 10). Our observations reveal a new cellular mechanism in malaria parasites that contributes to protein prenylation-dependent survival and sensitivity to thermal stress. Furthermore, our data suggest that compounds that target IPP synthesis or farnesylation may be more effective *in vivo* than *in vitro,* as parasites cycle through host temperature changes. As Hsp40 co-chaperone family members are amenable to small molecules inhibition^86, 87^, a combination therapy that targets both protein prenylation and heat shock proteins directly has the potential to be highly parasiticidal.

**Figure 10.**
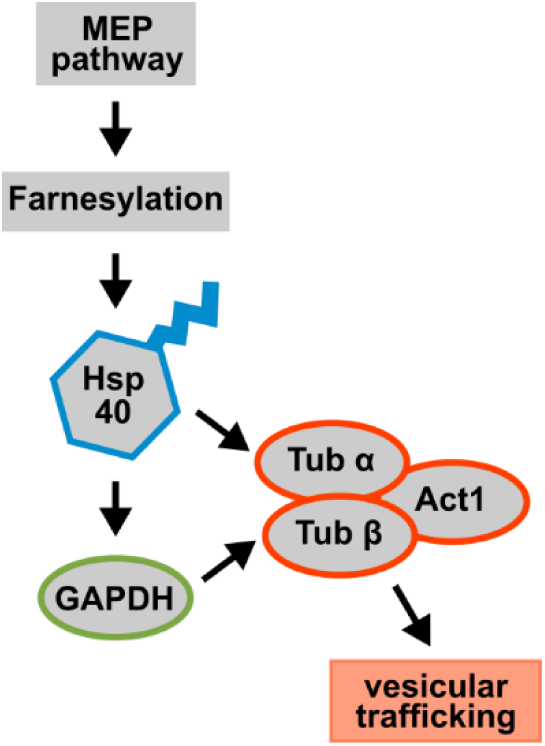
Model of farnesyl-HSP40 functional interactions with GAPDH and cytoskeleton components in the parasite. IPP-dependent HSP40 farnesylation promotes its interaction with Tubulin α chain (Tub α) Tubulin β chain (Tub β), Actin-1 (Act1), and GAPDH. ER-localized farnesyl-HSP40 likely mediates membrane trafficking in the early secretory pathway of the parasite, via direct interaction with these key client proteins.

## Materials and Methods

### Materials

All buffer components, salts, and enzyme substrates were purchased from Millipore Sigma (Burlington, MA), unless otherwise indicated.

### Statistical analysis

We plotted all data and performed all statistical analyses in GraphPad Prism software (version 8). All data are expressed as the mean ±SEM. For statistical analysis, we used 2-way ANOVA, t-test with Welch’s correction, and multiple unpaired t-test to compare results. To understand the interaction between treatment and temperature stress, we utilized a 2-way ANOVA and then adjusted p-values for multiple comparisons using Sidak’s multiple comparison test. For direct comparisons between control and treatment groups, we employed t-test with Welch’s correction, because standard deviations were not the same between groups. Mass spectrometry data was analyzed using multiple t-tests to efficiently compare results across conditions for each protein.

### Drug Inhibitors and Isoprenoids

Fosmidomycin (50mM; Millipore Sigma), FTI-277 (5mM; Tocris Bioscience, Bristol, UK), and IPP (30mM; Echelon Biosciences, Salt Lake City, UT) were each dissolved in water at concentrations indicated. GGTI-298 (20mM; Cayman Chemical, Ann Arbor, MI), BMS-388891 (20mM; kindly provided by Wesley Van Voorhis, U. Wash.), 2-bromopalmitate (100mM; Millipore Sigma), Farnesol (50mM; Millipore Sigma), and Geranylgeraniol (50mM; Echelon Biosciences) were each dissolved in 100% DMSO at concentrations indicated.

### Parasite strains and culture

Unless otherwise indicated, parasites were maintained at 37°C in 5% O_2_–5% CO_2_–90% N_2_ in a 2% suspension of human erythrocytes in RPMI medium modified with 27 mM NaHCO_3_, 11 mM glucose, 5 mM HEPES, 0.01 mM thymidine, 1 mM sodium pyruvate, 0.37 mM hypoxanthine, 10 µg/ml gentamicin, and 5 g/liter Albumax (Thermo Fisher Scientific, Waltham, MA). All experiments were conducted in wild-type strain 3D7 (MRA-102) obtained through the MR4 as part of the BEI Resources Repository, NIAID, NIH (www.beiresources.org).

### Heat and cold shock *P. falciparum* growth assays

Asynchronous cultures were diluted to 1% parasitemia. Cultures were treated with indicated drugs 24 hours prior to a 6-hour heat shock (40°C) or cold shock (25°C). Media (no drug) were exchanged post temperature shock. Cultures were split 1:6 after the collection of the Day 2 sample. IPP, F-OL, and GG-OL were supplemented in fresh media every second day. Samples were taken at indicated time points and fixed in phosphate-buffered saline (PBS)–4% paraformaldehyde. Cells were stained with 0.01 mg/ml acridine orange, and parasitemia was determined on a BD Biosciences LSRII flow cytometer (Thermo Fisher Scientific). All data represent means of results from ≥3 independent experiments using biological replicates.

### pCAM-BSD HSP40 plasmid construction

Functional genetic validation of HSP40 (PF3D7_1437900) was performed as previously described for PfDXR^20^ and PfIspD^45^. pCAM-BSD-HSP40^KO^ and pCAM-BSD-HSP40^ctrl^ vectors were derived from pCAM-BSD (gift from David Fidock, Columbia University), which includes a blasticidin-resistance cassette under transcriptional control by the *P. falciparum* D10 calmodulin 5′ UTR and the 3′ UTR of HRP2 from *P. berghei*. To construct pCAM-BSD-HSP40^KO^, the coding sequence for a segment of HSP40 near the N-terminus (bp 29 – 878) was inserted directly into 5′ of the *P. berghei* DHFR-thymidylate synthase 3′ UTR. This insert was constructed by PCR using primers (A) 5′-CCCGGGACTCTATGGGTGGTCAACAAG-3′ and (B) 5′-CTCGAGTCTCATGTAGGTCTTGGTTCC-3′ and restriction cloned using *Xma*I and *Xho*I sites. pCAM-BSD-HSP40^ctrl^ contained the coding sequence for the C-terminal end of HSP40 (bp 963 – 1619) and 237bp of the 3’ UTR, generated using (C) 5′-CCCGGGGAGGAACCAAGACCTACATGAG-3′ and (D) 5′-CTCGAGCATTTCACAGACACACACACAC-3′ as primers. This sequence was inserted at the same site, 5′ of the *P. berghei* DHFR-thymidylate synthase 3′ UTR. All constructs were verified by Sanger sequencing.

### Parasite transfections

Transfections were performed as previously described^45^. Briefly, 150 μg of plasmid DNA was precipitated and resuspended in Cytomix (25 mM HEPES [pH 7.6], 120 mM KCl, 0.15 mM CaCl_2_, 2 mM EGTA, 5 mM MgCl_2_, 10 mM K_2_HPO_4_). A ring-stage *P. falciparum* culture was washed with Cytomix and resuspended in the DNA/Cytomix solution. Cells were electroporated using a Bio-Rad Gene Pulser II electroporator at 950 µF and 0.31 kV. Electroporated cells were washed with media and returned to normal culture conditions. Parasites expressing the construct were selected by continuous treatment with 2 μg/mL Blasticidin S HCl (ThermoFisher Scientific). Transfectants were cloned by limiting dilution, and diagnostic PCRs were performed using genomic DNA from resultant transfectants using primer sets specific for episomal plasmids or genome integrants. Primer A and D sequences are given above: (X) 5′-TAAGAACATATTTATTAAACTGCAG-3′; (Y) 5′-GAAAAACGAACATTAAGCTGCCATA-3′.

### Southern Blots

Southern blots were used to assay the integration of either the pCAM-BSD-HSP40^ctrl^ or the pCAM-BSD-HSP40^KO^ plasmid. To assay the integration of pCAM-BSD-HSP40^KO^, genomic DNA was harvested from wild-type 3D7 *P. falciparum* and from pCAM-BSD-HSP40^KO^ transfectants #5 and #6 in Figure 1. These genomic DNA samples, along with pCAM-BSD-HSP40^KO^ plasmid, were digested with HpaII (New England Biolabs, Ipswich, MA). The knockout probe was prepared from PCR product generated using primers A and (HSP40_KO_R) 5’-ACTTCACTCACAATATCTTCACCTC-3’ (HSP40 bp 29–721) prior to southern blotting. To assay the integration of pCAM-BSD-PfHsp40^ctrl^, genomic DNA was harvested from wild-type 3D7 P. falciparum and from the continuously cultured pCAM-BSD-HSP40^ctrl^ transfectants #1, #2, #3, #4 from Figure 1. These genomic DNA samples, along with pCAM-BSD-HSP40^ctrl^ plasmid, were digested with SmlI (New England Biolabs). The control probe was prepared from PCR product generated using primers (HSP40_Ctrl_F) 5′-GAGGAACCAAGACCTACATGAG-3′ and (HSP40_Ctrl_R) 5′-ATGATCTTCATCGTCGTATGC-3′ (HSP40 bp 856–1574) prior to southern blotting.

### Generation of recombinant HSP40, HSP70, and GAPDH

An E. coli codon optimized HSP40 was produced (Genewiz, South Plainfield, NJ) and inserted via ligation-independent cloning into the isopropyl β-D-1-thiogalactopyranoside (IPTG) inducible BG1861 expression vector. This creates an N-terminal 6x-His tag fusion protein used for nickel purification. The expression plasmid was transformed into One Shot BL21(DE3)pLysS *E. coli* cells (Thermo Fisher Scientific). Overnight starter cultures were diluted 1:1000 and grown to an optical density (O.D.) of ∼0.6 where 1mM IPTG was added for 16 hours at 16°C. Cells were spun and stored at −80°C. Recombinant HSP70 (PF3D7_0818900) was expressed using the same conditions. In a similar manner, an E. coli codon optimized GAPDH (PF3D7_1462800) was produced with a few minor experimental differences. The GAPDH expression plasmid was transformed into One Shot BL21(DE3) E. coli cells (Thermo Fisher Scientific). 1mM IPTG was added for 2 hours at 37°C.

Expressed proteins were purified from cells using a sonication lysis buffer containing 1 mg/ml lysozyme, 20mM imidzazole, 1mM dithiothreitol, 1mM MgCl2, 10mM Tris HCl (pH7.5), 30 U benzonase, 1mM phenylmethylsulfonyl fluoride (PMSF), and cOmplete EDTA-free protease inhibitor tablets (Roche, Basel, Switzerland). Lysates were clarified using centrifugation and proteins were purified via nickel agarose beads (Gold Biotechnology, Olivette, MO), eluted with 300mM imidazole, 20mM Tris-HCl (pH 7.5) and 150mM NaCl. Eluted proteins were further purified via size exclusion chromatography using a HiLoad 16/60 Superdex 200 gel filtration column (GE Healthcare, Chicago, IL) using an AKTAExplorer 100 FPLC (GE Healthcare). Fast protein liquid chromatography buffer contained 100mM Tris-HCl (pH 7.5), 1mM MgCl2, 1mM DTT, and 10% w/v glycerol. HSP70 and GAPDH fractions containing purified protein were individually pooled, concentrated to ∼2mg/ml as determined via Pierce BCA Protein Assay Kit (Thermo Fisher Scientific), and stored by adding 50% glycerol for storage at −20°C. HSP40 fractions containing purified protein were pooled and further purified via anion exchange using a Mono Q anion exchange chromatography column (GE Healthcare) using an AKTAExplorer 100 FPLC (GE Healthcare). Anion exchange buffer contained 100mM Tris-HCl (pH 8.0), 1mM MgCl2, and 100mM NaCl. Purified fractions were concentrated to ∼2mg/ml as determined via Pierce BCA Protein Assay Kit (Thermo Fisher Scientific), glycerol was added to reach a concentration of 10% (wt/vol) and protein solutions were immediately flash frozen and stored at −80°C.

### HSP70 ATPase activity assays

Hydrolysis of ATP by HSP70 was measured using an EnzChek phosphate assay kit (Thermo Fisher Scientific). All reaction mixtures contained 50 mM ATP. HSP70 was added to the reaction ranging from 9 – 45 μg. HSP40 was used in reactions at 8.9 μg. Absorbance was measured every 12 seconds for 40 minutes. Slopes were calculated by using the Nonlinear Regression analysis tool in Prism (GraphPad Software). All data represent means of results from ≥3 independent experiments using biological replicates and performed with technical replicates.

### HSP40 and GAPDH antiserum generation

HSP40 and GAPDH rabbit polyclonal antisera were generated by Cocalico Biologicals (Reamstown, PA), using their standard protocol. Purified 6XHis-HSP40 or 6XHis-GAPDH were used as antigen and TiterMax was used as an adjuvant. Antiserum specificity was confirmed by immunoblotting of parasite lysate, RBCs, and purified protein. Further confirmation of anti-HSP40 specificity was conducted by performing IP analysis (discussed in section below).

### Immunofluorescence and Immuno-EM

For immunofluorescence labeling, infected RBCs at ∼8% parasitemia were fixed with 4% paraformaldehyde diluted in PBS. Fixed cells were washed with 50 mM ammonium chloride, permeabilized by treatment with 0.075% NP-40 in PBS and blocked using 2% bovine serum albumin in PBS. Cells were incubated with 1:5,000 rabbit polyclonal anti-HSP40 (described above). Hoechst 33258 (Thermo Fisher Scientific) was used as a nuclear counterstain. 1:1000 dilutions of Alexa Fluor 488 goat anti-rabbit IgG (Thermo Fisher Scientific, #A11008) were used as a secondary antibody. Images were obtained on an Olympus Fluoview FV1000 confocal microscope. For all immunofluorescence, minimal adjustments in brightness and contrast were applied equally to all images.

For immuno-EM, parasites were cultured at 2% hematocrit until reaching ∼6-8% parasitemia. Parasites were magnetically sorted from uninfected RBCs and ring stage parasites via MACS LD separation columns (Miltenyi Biotech, Bergisch Gladbach, Germany). Parasites were collected by centrifugation and fixed for 1 hour on ice in 4% paraformaldehyde in 100 mM PIPES/0.5 mM MgCl2, pH 7.2. Samples were then embedded in 10% gelatin and infiltrated overnight with 2.3 M sucrose/20% polyvinylpyrrolidone in PIPES/MgCl2 at 4°C. Samples were frozen in liquid nitrogen and then sectioned with a Leica Ultracut UCT7 cryo-ultramicrotome (Leica Microsystems, Wetzlar, Germany). 50 nm sections were blocked with 5% fetal bovine serum/5% normal goat serum for 30 min and subsequently incubated with primary antibody for 1 hour at room temperature. Primary antibodies used include anti-HSP40 (1:250) and anti-PDI (1D3) mouse 1:100 (ADI-SPA-891-D; Enzo Life Sciences, Farmingdale, NY). Secondary antibodies were added at 1:30 for 1 hour at room temperature. Secondary antibodies included 12 nm Colloidal Gold AffiniPure Goat anti-Mouse IgG (H+L)(115-205-146; Jackson ImmunoResearch, West Grove, PA) and 18 nm Colloidal Gold AffiniPure Goat anti-Rabbit IgG (H+L)(111-215-144; Jackson ImmunoResearch). Sections were then stained with 0.3% uranyl acetate/2% methyl cellulose, and viewed on a JEOL 1200 EX transmission electron microscope (JEOL USA Inc., Peabody, MA) equipped with an AMT 8 megapixel digital camera and AMT Image Capture Engine V602 software (Advanced Microscopy Techniques, Woburn, MA). All labeling experiments were conducted in parallel with controls omitting the primary antibody. Quantification of membrane associated HSP40 was performed in micrographs where both a nucleus and food vacuole were present. Images were blinded and scored for total number labeled HSP40 and membrane-associated HSP40.

### Membrane fraction preparation and immunoblotting

Asynchronous parasites were released from RBCs with 0.1% saponin, washed in cold PBS and resuspended in 100 – 300 μl DI-water with 1mM PMSF and cOmplete EDTA-free protease inhibitor tablet (Roche). Resuspended pellets were freeze-thawed three times with liquid nitrogen/37°C water bath. A total lysate sample was taken at this point in the protocol. The membranes were pelleted (14,000 RPM, 30 minutes, 4°C) and the supernatant was collected as the soluble fraction. Pellets were washed once with ice-cold PBS and pelleted (as before) before resuspending pellets in 100 – 300 μl (depending on sample amount) RIPA buffer (Cell Signaling Technology, Danvers, MA) containing 1% CHAPS and 1% ASB-14. Samples were sonicated three times with a microtip and incubated at 42°C with shaking at 800 RPM for 45 min. The samples were then centrifuged (14,000 RPM, 30 minutes, 4°C) and the resulting supernatant was collected as the membrane fraction. 4X SDS sample buffer was added, samples were boiled for 10 minutes and loaded on 4–20% Mini-PROTEAN TGX gradient gels (Bio-Rad Laboratories, Hercules, CA).

For immunoblotting, proteins were transferred onto PVDF using wet transfer with 20% methanol. Blots were blocked either 1 hour at 25°C or overnight at 4°C with 2% bovine serum albumin– 0.1% Tween 20–PBS. Primary antibodies were used at the following dilutions: 1:5,000 rabbit anti-HSP40, 1:5,000 rabbit anti-GAPDH, 1:10,000 rabbit anti-HAD1^88^, 1:5,000 rabbit anti-Hsp70 (AS08 371; Agrisera Antibodies, Vännäs, Sweden), 1:500 mouse anti-PM-V^89^, and 1:5,000 mouse anti-EXP-2 clone 7.7^90^. For all blots, 1:20,000 horseradish peroxidase (HRP)-conjugated goat anti-rabbit IgG antibody (Thermo Fisher Scientific, 65-6120) or 1:5,000 HRP-conjugated goat anti-mouse IgG antibody (Thermo Fisher Scientific, G-21040) were used as secondary antibodies. Anti-HAD1 or anti-Hsp70 and anti-PM-V or anti-Exp-2 were used as loading controls from total lysate and membrane fractions, respectively. All blot images are representative of results of a minimum of 3 independent experiments using biological replicates.

### Immunoprecipitation of HSP40 from parasite lysate and protein mass-spectrometry

Pre-inoculated rabbit antisera and anti-HSP40 were coupled to magnetic beads using the DynabeadsTM Antibody Coupling Kit as per the manufacturers protocol (Thermo Fisher Scientific, 14311D). Parasite pellets were harvested from RBCs using 0.1% saponin. Isolated parasite pellets were lysed in 300μl of buffer containing 25mM Tris-HCl (pH 7.5), 100mM NaCl, 5mM EDTA, 0.5% Triton-X 100, and one cOmplete Mini EDTA-free protease inhibitor tablets (Roche) (per 10mL of buffer). Resuspended pellets were homogenized using a LabGEN homogenizer (Cole-Parmer, Vernon Hills, IL) in three rounds of 30-second intervals with 60 seconds of rest on ice. Lysate was centrifuged at 14,000 RPM for 10 minutes at 4°C. The resulting soluble lysate was diluted by adding 450μl of binding buffer containing 25mM Tris-HCl (pH 7.5) and 150mM NaCl. Diluted lysate was added to antisera coupled beads that were previously washed three times with the same binding buffer. Lysate and beads were rotated for 2 hours at 4°C. Beads were then washed three times with wash buffer (25mM Tris-HCl (pH 7.5) and 500mM NaCl) and flow through was discarded. Immunoprecipitated proteins were eluted using elution buffer with 200mM glycine (pH 2.5) for 30 seconds and eluted sample was neutralized with 1M Tris-HCl (pH 7.5). Samples were then flash frozen and stored at −80° for immunoprecipitate identification via protein mass-spectrometry.

Immunoprecipitates were identified via protein mass-spectrometry by the Proteomics and Mass Spectrometry Core at the Donald Danforth Plant Science Center (St. Louis, MO). Stored samples were submitted in solution for protein mass-spectrometry. Total spectra were blasted to the *Plasmodium falciparum* (3D7) proteome UniProt UP000001450 and analyzed in Scaffold (Proteome Software). The Table S1 protein list for untreated control was generated by identifying proteins that had an average of greater than 10 peptide count and a fold change >1.4 when comparing peptides found in anti-HSP40 versus pre-inoculated rabbit antisera. FSM-treated and FTI-treated fold change was calculated by first normalizing peptide counts for each protein in each sample to the HSP40 peptide number. Then fold change for each protein was calculated by dividing the average in the drug-treated sample by the average in untreated controls. p-values were calculated using unpaired t-test not assuming consistent SD and were not corrected for multiple comparisons. The biological process gene ontology analysis (Fig. 8A) was done using PANTHER (pantherdb.org). Table S1 proteins were used as the cohort for untreated control. The FSM-treated and FTI-treated protein lists used for gene ontology analysis were generated using the same criteria used to generate Table S1 (>10 average peptide count and fold change >1.4). Volcano plots (Fig. 8B,C) were calculated by first normalizing peptide counts for each protein in each sample to the HSP40 peptide number. Fold change for each protein was calculated by dividing the average in the drug-treated sample by the average in untreated controls and p-values were determined by performing multiple unpaired t-tests not assuming consistent SD and not corrected for multiple comparisons.

### Metabolite profiling

60mL of sorbitol-synchronized early trophozoites cultured at 4% hematocrit until reaching ∼7-11% parasitemia were isolated using 0.1% saponin, washed with ice-cold PBS, and frozen at −80°C. Glycolysis and pentose phosphate pathway intermediates were extracted via the addition of glass beads (212-300 u) and 600 µL chilled H2O:chloroform:methanol (3:5:12 v/v) spiked with PIPES (piperazine-N,N′-bis(2-ethanesulfonic acid)) as internal standard. The cells were disrupted with the TissueLyser II instrument (Qiagen, Hilden, Germany) using a microcentrifuge tubes adaptor set pre-chilled for 2 min at 20 Hz. The samples were then centrifuged at 16,000 g at 4 °C, the supernatants collected, and pellet extraction repeated once more. The supernatants were pooled and 300 µL chloroform and 450 µL of chilled water were added to the supernatants. The tubes were vortexed and centrifuged. The upper layer was transferred to a new tube and dried using a speed-vac. The pellets were re-dissolved in 100 µL of 50% acetonitrile.

For liquid chromatography (LC) separation of the glycolysis/pentose phosphate pathway intermediates, an InfinityLab Poroshell 120 HILIC (2.7 um, 150 x 2.1 mm, Agilent) was used flowing at 0.5 mL/min. The gradient of the mobile phases A (20 mM ammonium acetate, pH 9.8, 5% ACN) and B (100% acetonitrile) was as follows: 85% B for 1 min, to 40% B in 9 min, hold at 40% B for 2 min, then back to 85% B in 0.5 min. The LC system was interfaced with a Sciex QTRAP 6500+ mass spectrometer equipped with a TurboIonSpray (TIS) electrospray ion source. Analyst software (version 1.6.3) was used to control sample acquisition and data analysis. The QTRAP 6500+ mass spectrometer was tuned and calibrated according to the manufacturer’s recommendations. Metabolites were detected using MRM transitions that were previously optimized using standards. The instrument was set-up to acquire in negative mode. For quantification, an external standard curve was prepared using a series of standard samples containing different concentrations of metabolites and fixed concentration of the internal standard. The limit of detection for glycolysis and pentose phosphate pathway intermediates were as follows: glucose 6-phosphate and glucose 1-phosphate/fructose 6-phosphate 0.5 µM; glyceraldehyde 3-phosphate and ribulose 5-phosphate 1 µM; erythrose-4-phosphate 1.5 µM; pyruvate, 2/3-phosphoglycerate, phosphoenolpyruvate, and sedoheptulose-7-phosphate 2 µM; fructose 1,6-bisphosphate 3.9 µM. Resulting metabolite levels were normalized to parasitemia levels for each individual sample and reported as attogram per cell (ag/cell). Levels were averaged between three biological replicates and compared between control, FSM (5 µM for 24 hours), and FTI (10 µM for 24 hours).

## Acknowledgments.

We thank Daniel Goldberg (Washington University) for supplying the anti-Exp-2 and anti-PM-V antibodies and Wesley Van Voorhis (University of Washington) for supplying BMS-388891. We thank Wandy Beatty (Washington University) for assistance with immuno-electron microscopy and Sophie Alvarez (University of Nebraska-Lincoln) for assistance with metabolomics profiling. Research reported in this publication was supported by NIH/NIAID AI103280 (AOJ), AI123433 (AOJ), AI130584 (AOJ), AI144472 (AOJ), S10 RR026891, T32AI007172-37 (ESM), 1F32AI138373-01 (ESM), and T32GM007067 (AJJ). Dr. John is an Investigator in the Pathogenesis of Infectious Diseases of the Burroughs Welcome Fund. Support is acknowledged from the National Science Foundation under Grant No. DBI-0922879 for acquisition of the LTQ-Velos Pro Orbitrap LC-MS/MS (Danforth Plant Sciences Center). We are grateful for the discussions and input from both the Odom John and Goldberg laboratory members.

